# Enhancing the Antiviral Efficacy of RNA-Dependent RNA Polymerase Inhibition by Combination with Modulators of Pyrimidine Metabolism

**DOI:** 10.1101/2020.03.24.992230

**Authors:** Qi Liu, Amita Gupta, Ayse Okesli-Armlovich, Wenjie Qiao, Curt R. Fischer, Mark Smith, Jan E. Carette, Michael C. Bassik, Chaitan Khosla

## Abstract

Genome-wide analysis of the mode of action of GSK983, a potent antiviral agent, led to the identification of dihydroorotate dehydrogenase (DHODH) as its target, along with the discovery that knockdown of genes in pyrimidine salvage pathways sensitized cells to GSK983. Because GSK983 is an ineffective antiviral in the presence of physiological uridine concentrations, we explored combining GSK983 with pyrimidine salvage inhibitors. We synthesized and evaluated analogs of cyclopentenyl uracil (CPU), an inhibitor of uridine salvage. We found that CPU was efficiently converted into its triphosphates in cells. When combined with GSK983, it led to large drops in cellular UTP and CTP pools. Consequently, CPU-GSK983 suppressed dengue virus replication in the presence of physiological concentrations of uridine. In addition, the CPU-GSK983 combination markedly enhanced the effect of RNA-dependent RNA polymerase (RdRp) inhibition on viral genome infection. Our findings highlight a new host-targeting strategy for potentiating the antiviral activities of RdRp inhibitors.

## INTRODUCTION

Significant progress in the development of antiviral drugs has come by targeting viral proteins with small molecules (Jordheim et al., 2013; Lou et al., 2014). For examples, compounds like aciclovir and zidovudine block viral reverse transcriptase to treat herpes simplex virus and HIV infections, respectively, and RNA-dependent RNA polymerase (RdRp) inhibitors like dasabuvir and sofosbuvir are used to treat hepatitis C virus infections (Nováková et al., 2018). More recently, the broad-spectrum RdRp inhibitor remdesivir (Gordon et al., 2020) has entered clinical trials for coronavirus disease 2019 (Covid-19). Meanwhile, targeting host proteins required for viral propagation is emerging as an attractive alternative that may circumvent the emergence of resistance (Bekerman and Einav, 2015). For example, maraviroc inhibits the human chemokine receptor CCR5, and is therefore used to treat multidrug-resistant HIV (Lieberman-Blum et al., 2008). Additionally, pyrimidine biosynthesis has emerged as a potential host-targeting strategy for antivirals (Okesli et al., 2017). Here, we focus on devising a host-targeting antiviral approach for the treatment of RNA viruses, which cause many serious diseases such as hepatitis, influenza, Ebola, dengue, and Covid-19.

In mammalian cells, pyrimidine biosynthesis is a tightly regulated metabolic process. Two complementary pathways – *de novo* biosynthesis and pyrimidine salvage – are responsible for producing UTP and CTP for host as well as viral RNA synthesis (Figure 1). *De novo* pyrimidine biosynthesis is a resource-intensive process. In contrast, salvage occurs via phosphorylation of UMP and CMP derived from intracellular RNA degradation or via facilitated transport and phosphorylation of extracellular uridine, whose plasma concentration is tightly controlled in the low micromolar range (Traut, 1994). Recently we discovered that GSK983, a broad-spectrum antiviral agent first reported in 2009 (Harvey et al., 2009), is a potent inhibitor of dihydroorotate dehydrogenase (DHODH), a rate-limiting step in *de novo* pyrimidine biosynthesis (Deans et al., 2016). In the course of those unbiased genome-wide studies, we also found that knockdown of uridine/cytidine kinase 2 (UCK2) and cytidine monophosphate kinase 1 (CMPK1) in the pyrimidine salvage pathway strongly sensitized cells to growth inhibition by GSK983(Deans et al., 2016). This finding was consistent with the observation that GSK983 often lacks antiviral efficacy *in vivo* despite high potency *in vitro* presumably due to salvage metabolism of circulating uridine by virus-infected cells (Traut, 1994; Wang et al., 2011).

**Figure 1.**
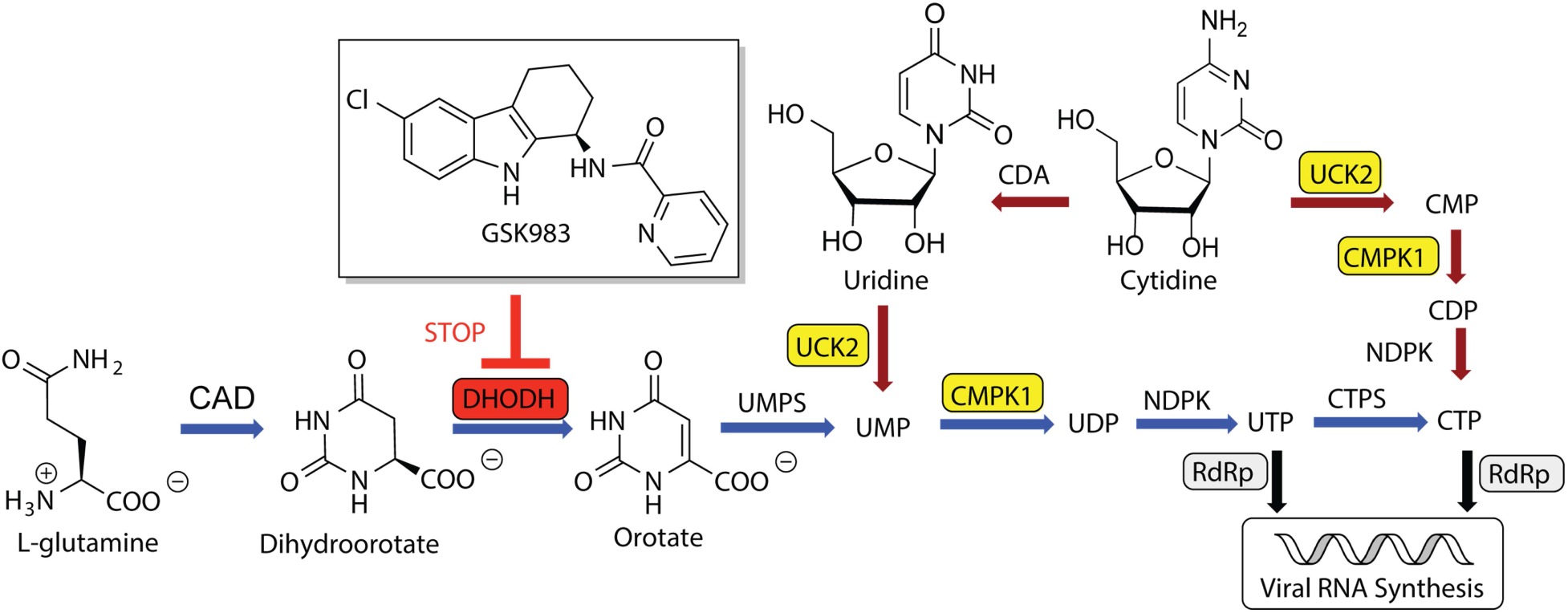
*De novo* and salvage biosynthesis of pyrimidine nucleotides for host and viral RNA synthesis. GSK983 is a DHODH inhibitor. Genes that sensitize cells to GSK983 are highlighted in yellow boxes. Reactions shown with blue arrows comprise the *de novo* biosynthetic pathway, whereas those with red arrows comprise the salvage pathway.

To restore the antiviral efficacy of GSK983 in the presence of extracellular uridine, we therefore sought to inhibit pyrimidine salvage. Cyclopentenyl uridine (CPU) is a carbocyclic analogue of uridine that has been shown to inhibit human UCK2 (Lim et al., 1984). To our surprise, we learned that the antiviral activity of CPU is due to its remarkable ability to deplete intracellular pyrimidine nucleotide pools via salvage biosynthesis pathways. Our findings led us to redirect our search for a fundamentally new type of combination chemotherapy for RNA viruses, as described below.

## RESULTS

### Diversity-oriented Syntheses of CPU Analogs

Our search for lead inhibitors of pyrimidine salvage was inspired by earlier reports on the biological activity of CPU and cyclopentenyl cytosine (CPC), which were shown to block uridine salvage *in vitro* and *in vivo* (Cysyk et al., 1995; Lim et al., 1984; Moyer et al., 1985; Schimmel et al., 2007). In these and other reports, CPU was found to be remarkably well-tolerated, whereas CPC was considerably more cytotoxic (Blaney et al., 1992; Ford et al., 1991; Song et al., 2001). This contrast is likely due to downstream inhibition of the CTP synthetase by CPC(Schimmel et al., 2007).

Due to the high toxicity of CPC, we undertook structure-activity relationship (SAR) analysis of CPU wherein uracil nucleobase was maintained intact or only modified at the C5 position. Meanwhile, the C-5’ substituent of CPU was modified, because this is the site of UCK2-catalyzed phosphorylation. In order to rapidly access both nucleobase and carbocyclic moiety analogs, we implemented a diversity-oriented synthetic approach featuring a Mitsunobu reaction as the strategic transformation (Choi et al., 2012).

We first synthesized CPU analogs with nucleobase modifications (Figure 2*A*). Mitsunobu reactions between the common cyclopentenyl moiety (**1**) (Choi et al., 2004) and the benzoyl protected uracil, C(5)-fluoro-uracil, C(5)-iodo-uracil or thymine (**2a**-**2d**) (Racine et al., 2014) furnished the carbon skeleton of C-5 analogs. Removal of acetal, benzoyl and TBDPS groups then delivered analogs **4a**-**4d**.

**Figure 2.**
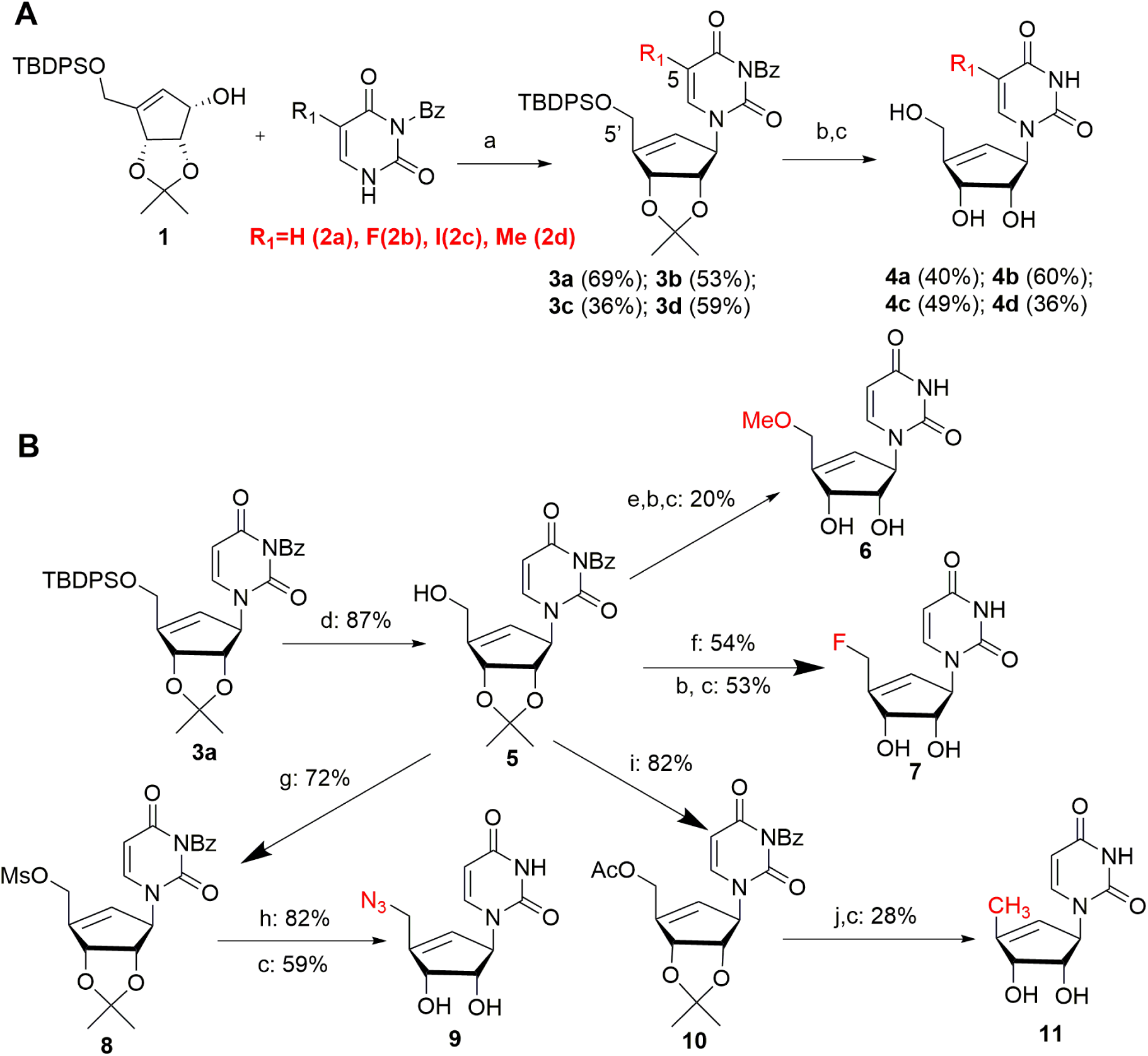
Diversity-oriented syntheses of CPU analogs. (*A*) Modification of the C5 position to afford uracil moiety analogs. (*B*) Modification of the C5’ position to furnish carbocyclic moiety analogs. a) Diethyl azodicarboxylate (DEAD), PPh_3_, THF; b) NH_3_, MeOH; c) HCl, THF; d) TBAF, THF; e) Ag_2_O, MeI, acetone; f) DAST, CH_2_Cl_2_; g) MsCl, Et_3_N, CH_2_Cl_2_; h) NaN_3_, DMF; i) AcCl, CH_2_Cl_2_; j) Pd(OH)_2_, cyclohexene, ethanol. TBDPS=*tert*-Butyldiphenyl silyl; Bz=Benzoyl; TBAF=Tetra-*n*-butylammonium fluoride, DAST=Diethylaminosulfur trifluoride, Ms= Methanesulfonyl, Ac=acetyl.

Next, we turned to synthesis of CPU analogs modified at the C5’ position of the cyclopentenyl moiety (Figure 2*B*). Selective deprotection of TBDPS with TBAF revealed the primary hydroxyl group (Song et al., 2001), which was then methylated in the presence of Ag_2_O (Cosme G. Francisco et al., 2001) to afford the protected 5’-methoxy-CPU. Alternatively, the alcohol could be converted to a terminal fluoride with diethylamino sulfurotrifluoride (DAST) (Moon et al., 2004), resulting in protected 5’-fluoro CPU. Mesylation or acetylation of **5**, followed by NaN_3_ substitution (Schaudt and Blechert, 2003) or Pd(OH)_2_/C catalyzed hydrogenation (Bianco et al., 1989) afforded the protected 5’-azido- and 5’-deoxy-CPU, respectively. The resulting protected intermediates were treated with methanolic ammonia and/or HCl to provide analogs **6**,**7, 9** and **11**.

### Enzymatic Analysis of CPU Analogs Against UCK2 and CMPK1

In order to guide our SAR studies, we developed an enzymatic assay to evaluate the inhibitory effect of each CPU analog against recombinant human UCK2, which was expressed and purified in *E. coli*. A continuous assay system was optimized by coupling UCK2 activity to the pyruvate kinase (PK) reaction, which in turn was coupled to lactate dehydrogenase (LDH) (Tomoike et al., 2017). Overall reaction progress was continuously monitored by detecting the UV absorption change at 340 nm, which was correlated with the ATP consumption by UCK2.

The effect of 250 μM of each CPU analog on UCK2 activity was evaluated in the presence of 50 μM uridine. CPU and 5-F-CPU showed much higher ATP consumption compared to other analogs (Figure 3*A*), suggesting these two compounds were substrates of UCK2. We then individually measured the steady state kinetic parameters of UCK2 using uridine, CPU and 5-F-CPU as substrates. Their *K*_*M*_ values were 86, 25, and 41 μM, and their *k*_*cat*_ */ K*_*M*_ values were 2.6×10^5^, 1.5×10^5^, and 7.3×10^4^ s^−1^×M^−1^ (Figure 3*B*). As predicted by the relative magnitude of these kinetic parameters, addition of CPU or 5-F-CPU to an assay mixture containing UCK2, uridine and ATP resulted in a dose-dependent decrease in the rate of UMP synthesis (Figure 3*C*).

**Figure 3.**
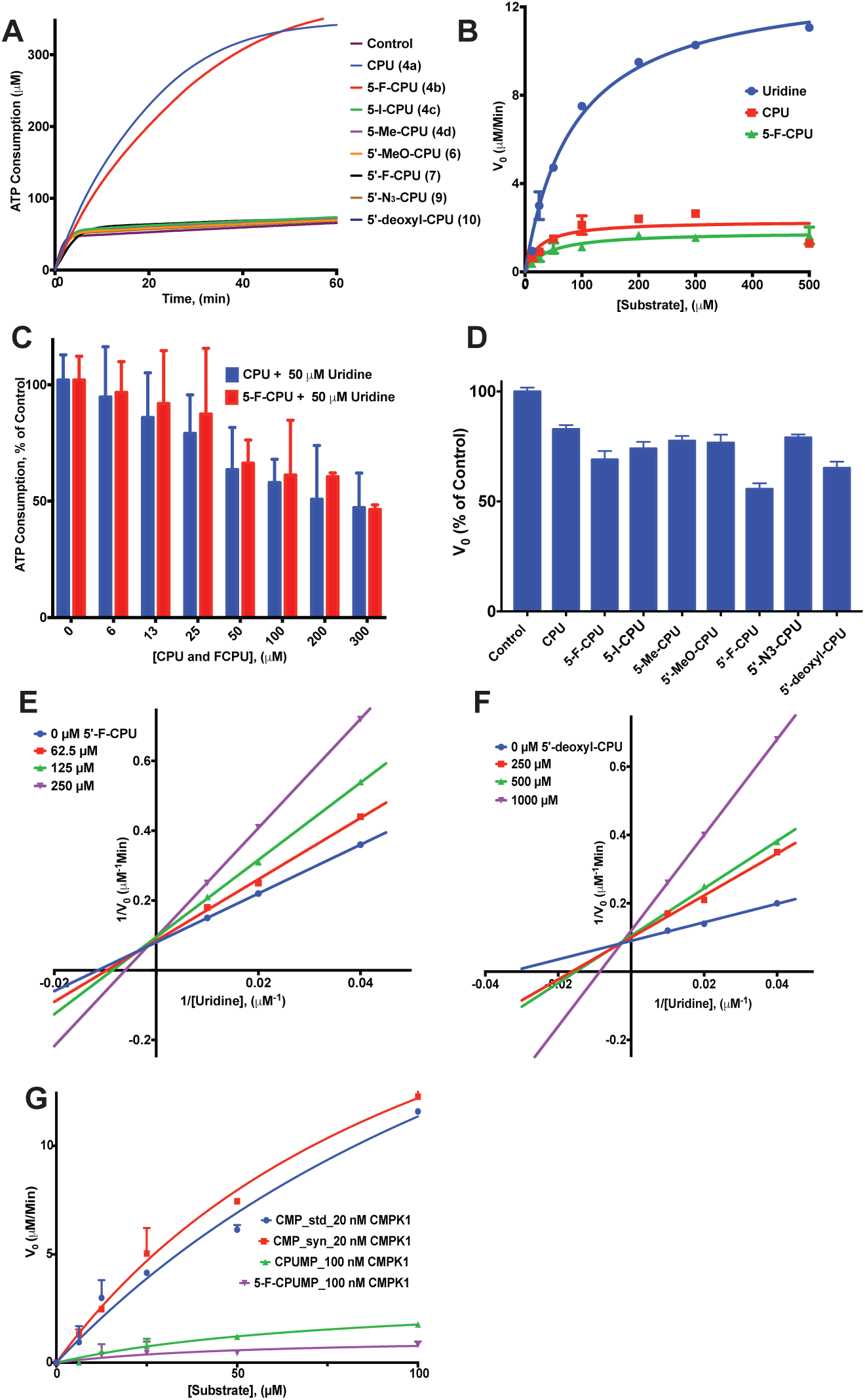
Enzymatic analysis of CPU analogs. (*A*) CPU and 5-F-CPU are UCK2 substrates, as observed by addition of 250 μM of each CPU analogue to a UCK2 reaction mixture that also contains 50 μM uridine. All other analogs tested are not UCK2 substrates. (*B*) Steady-state kinetic analysis of uridine, CPU and 5-F-CPU as UCK2 substrates. Error bars represent ± S.D. of three replicates. (*C*) Comparative ATP consumption by UCK2 in the presence of 50 μM uridine and varying concentrations of CPU (blue bars) or 5-F-CPU (red bars). Error bars represent ± S.D. of three replicates. (*D*) Initial velocities calculated from the data shown in panel (a). (*E*) and (*F*) Lineweaver-Burk analysis of 5’-F-CPU and 5’-deoxy-CPU as UCK2 inhibitors. (*G*) Comparative activity of CMPK1 on CMP, CPU-MP and 5-F-CPU-MP. An authentic standard of CMP (std) was tested alongside a UCK-synthesized sample of the same compound (syn). Error bars represent ± S.D. of three replicates.

While CPU and 5-F-CPU were the only UCK2 substrates identified from our panel of carbocyclic nucleoside analogs, some of the other agents had measurable inhibitory activity against this enzyme (Figure 3*D*). The *Ki* values of the two most potent competitive inhibitors, 5’-F-CPU and 5’-deoxy-CPU, were 170 μM and 230 μM, respectively (Figures 3*E* and 3*F*).

Given that CPU and 5-F-CPU could be phosphorylated by UCK2, we sought to establish whether the corresponding monophosphates were substrates of human CMPK1. For this purpose, human CMPK1 was expressed in *E. coli* and purified. We then employed recombinant UCK2 to synthesize CMP, CPU-MP and 5-F-CPU-MP and confirmed their identities by LC-MS/MS (Figure S1). The results shown in Figure 3*G* demonstrate that both CPU-MP and 5-F-CPU-MP are substrates of CMPK1, but that the former is a better substrate. Identities of their corresponding products, CPU-DP and 5-F-CPU-DP, were also confirmed by LC-MS/MS (Figure S2).

### LC-MS Analysis of Intracellular Nucleotides

To understand the metabolic implications of the above biochemical findings, we developed a LC-MS/MS based assay that facilitated measurement of the effects of CPU,5-F-CPU and 5’-F-CPU on intracellular pyrimidine nucleotide levels. To minimize the perturbative effects of sample quenching and analysis on the physiological concentrations of these metabolites, we adapted an earlier protocol for growing cells on glass cover slips to facilitate rapid washing (Figure S3) (Martano et al., 2014). LC-MS of nucleotides (Table S1 for MS/MS transitions of selected precursors) was performed using a dynamic multiple reaction monitoring (dMRM) method (Sartain et al, 2016). Addition of micromolar concentrations of medronic acid into the mobile phase markedly increased the sensitivity for detecting di- and tri-nucleotides by this method (Hsiao et al., 2018).

With optimized sampling and analysis method in hand, pyrimidine nucleotide levels were measured in cells cultured with either CPU, 5-F-CPU or 5’-F-CPU in combination with 1 μM GSK983 and 5 μM uridine for 6 h (Figure 4*A*). Inclusion of CPU or 5-F-CPU led to large decreases in UTP and CTP levels compared to cells treated with GSK983 alone; these effects grew more pronounced at 12 h (Figure S4). CPU depleted UTP and CTP concentrations more strongly than 5-F-CPU. Triphosphates of both CPU and 5-F-CPU could be detected by LC-MS (Figure 4*B*), suggesting that nucleoside diphosphate kinase, a mammalian enzyme known to have broad substrate scope (Lascu and Gonin, 2000), could convert CPU-DP and 5-F-CPU-DP into their corresponding nucleoside triphosphate analogs. Inclusion of CPU or 5-F-CPU also led to moderate decreases in UMP, CMP, UDP, and CDP levels compared to cells treated with GSK983 alone at both 6 h and 12 h (Figure S4).

**Figure 4.**
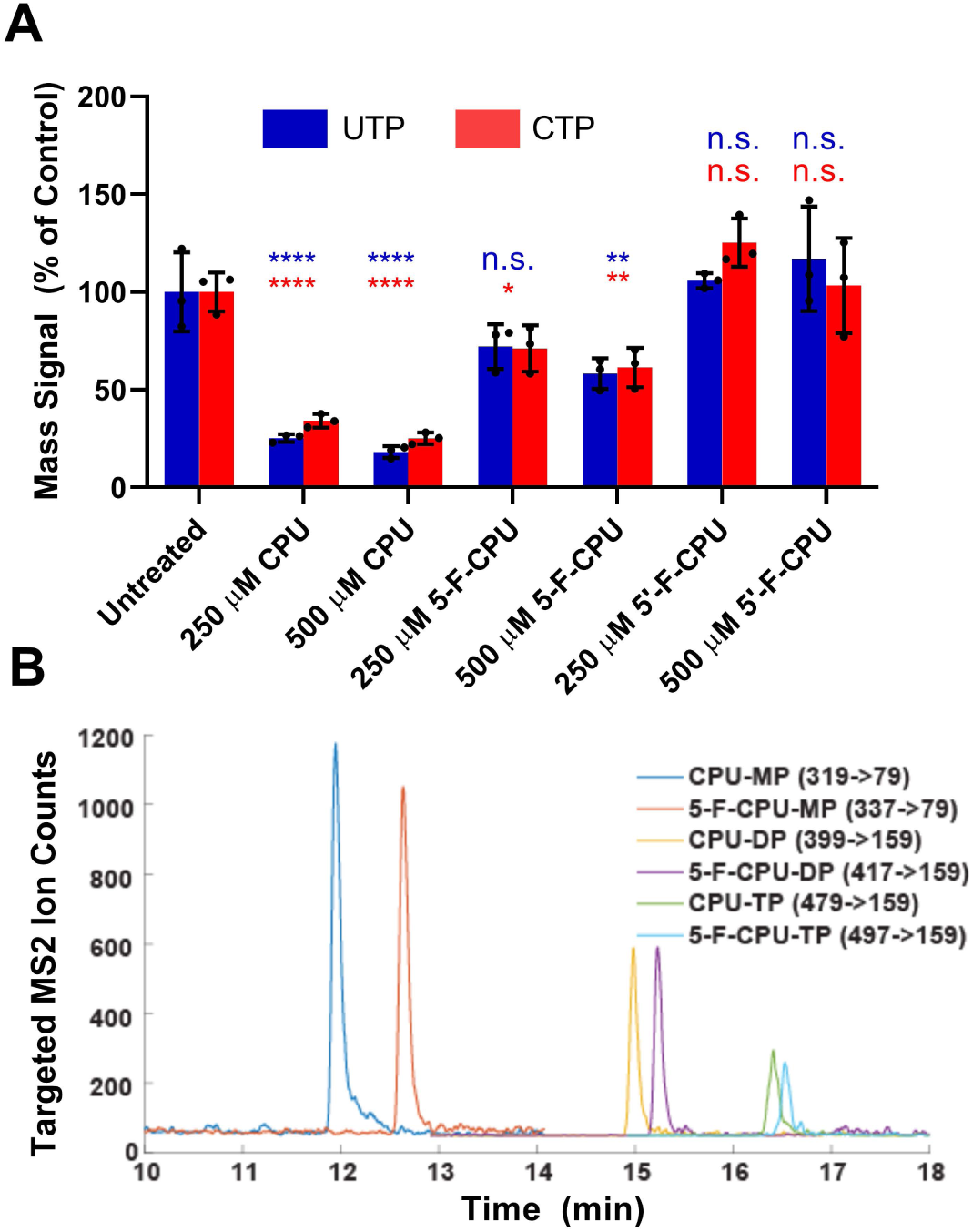
LC-MS analysis of intracellular nucleotides. (*A*) LC-MS analysis of intracellular uridine and cytidine nucleotide levels after 6 h treatments. In all assays, the culture medium was supplemented with 5 μM uridine and 1 μM GSK983. Error bars represent ± S.D. of three biological replicates with statistical comparisons to the untreated control provided for UTP levels (blue) and CTP levels (red) as not significant (n.s), * (*p* <0.05), ** (*p* < 0.01), *\*\*\** (*p* < 0.001) and **** (*p* < 0.0001) as calculated by one way ANOVA, Dunnett’s test. (*B*) LC-MS detection of mono-, di- and tri-phosphates of CPU and 5-F-CPU in cells.

Despite its UCK2 inhibitory activity *in vitro*, 5’-F-CPU did not deplete intracellular nucleotide levels and even appeared to increase levels of some intracellular nucleotides (Figure 4A and Figure S4). This contrast suggests that the activity of CPU and 5-F-CPU is derived primarily from their flux through pyrimidine metabolism pathways and not from their inhibition of UCK2.

### Antiviral and Cytotoxic Activities of Selected CPU Analogs

From above enzymological and metabolic data, we hypothesized that the combination of CPU and GSK983 would be more effective at inhibiting dengue virus replication than any other combination. Using an infectious clone of dengue serotype 2 (DENV-2) strain 16681 engineered to express a luciferase reporter (Deans et al., 2016; Hierholzer and Killington, 1996; Marceau et al., 2016), the efficacy and cytotoxicity of each combination treatment was tested in infected or uninfected cultures of the A549 lung carcinoma cell line (Figure 5*A*). In all assays, culture medium was supplemented with 20 μM uridine to mimic physiological plasma concentrations. In the presence of 20 µM exogenous uridine, 1 µM of the *de novo* pyrimidine synthesis inhibitor GSK983 alone did not significantly inhibit viral infection or cellular viability (Figure 5*A*). In line with our original hypothesis, combination of GSK983 with either 500 µM CPU and 5-F-CPU caused significant reductions in viral infection unlike the other cyclopentenyl compounds tested (Figure 5*A*). However, only CPU appeared to have a workable therapeutic index at this concentration and time point.

**Figure 5.**
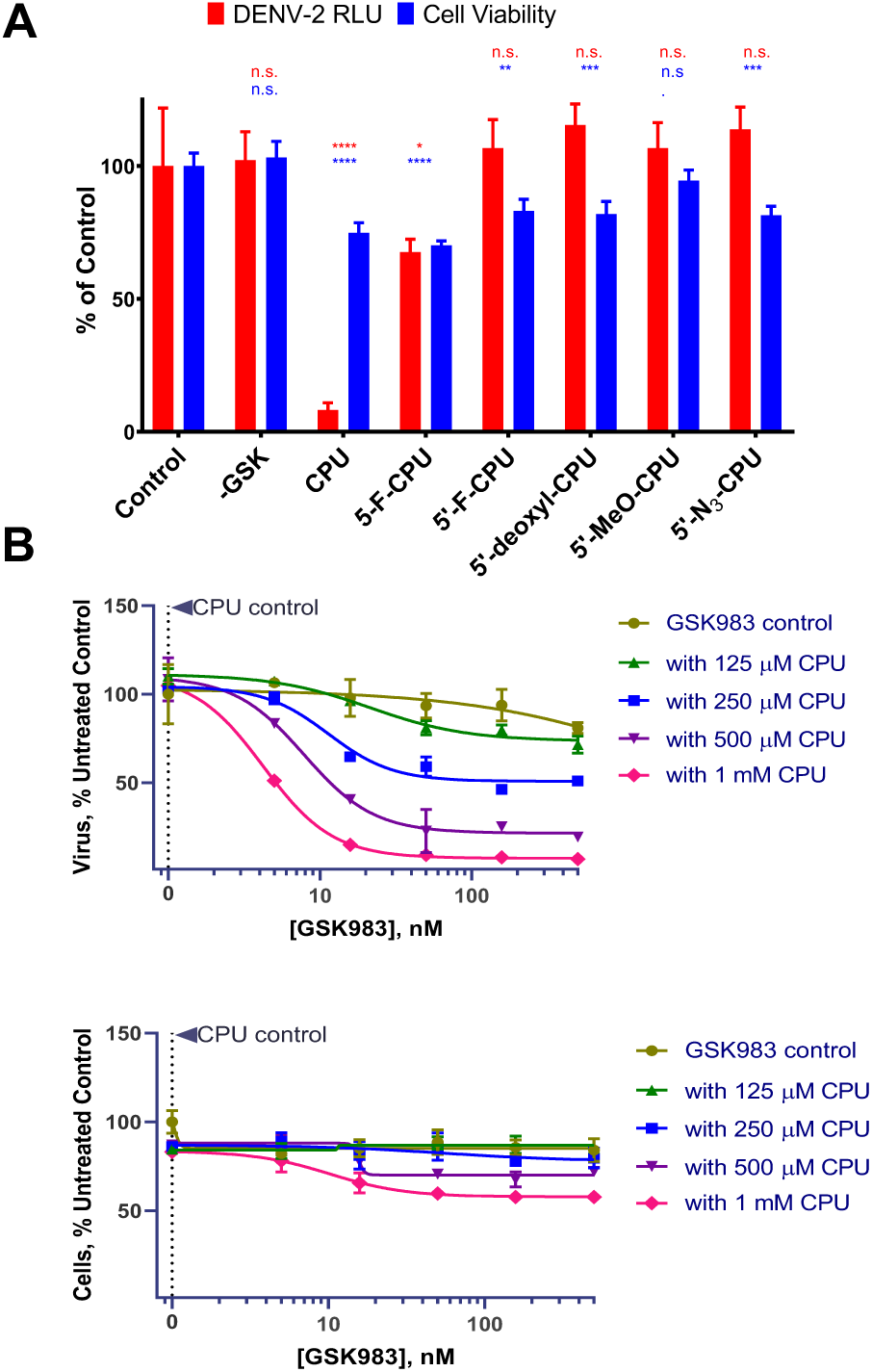
Antiviral and cytotoxic activities of modulators of pyrimidine metabolism in the presence of exogenous uridine. *(A)* Effect of 1 μM GSK983 and 500 μM of individual CPU analogs on the viability of A549 cells (blue) and on replication of luciferase-expressing DENV-2 virus (red) at 72 h. Error bars represent ± S.D. of three replicates. *(B)* Effects of CPU-GSK983 combination therapy on dengue virus replication and cell proliferation in the presence of exogenous uridine at 48 h. Error bars represent ± S.D. of two replicates.

To further examine the efficacy and cytotoxicity of CPU, we conducted a more comprehensive checkerboard analysis of the CPU-GSK983 combination. While neither GSK983 nor CPU were effective as single agents in physiological uridine levels, the combination of both molecules successfully suppressed dengue virus infection (Figure 5*B*). For example, combining 0.2 μM GSK983 and 250 μM CPU inhibited ∼ 50% of virus replication (Figure 5*B*). At a CPU dose of 1 mM, virus replication was suppressed almost completely. Notably, the combination treatment had much less effect on A549 cell growth, suggesting that combinations of GSK983 and CPU could be selective for inhibition of virus but not host replication.

To further demonstrate that intracellular nucleotide depletion is correlated with antiviral activity, we also tested the effects of combining GSK983 with a broad inhibitor of nucleoside transport, dipyridamole (DPY). This drug significantly suppressed intracellular nucleotide levels (Figure S4) and consequently inhibited viral replication (Figure S5).

### Improving the Therapeutic Index of RdRp Inhibitor R1479 By Combination Treatment Targeting Pyrimidine Biosynthesis

Given that the CPU-GSK983 combination was able to selectively inhibit viral infection through modulation of intracellular nucleotide pools, we hypothesized that the combination should potentiate the effects of an RdRp inhibitor on viral genome replication. Among RdRp inhibitors, the nucleoside inhibitor (NI) class is the largest (Klumpp et al., 2006) and has progressed the furthest in studies against dengue virus (Lim et al., 2015). Many NIs are nucleoside analogs that rely on intracellular phosphorylation to form triphosphate substrates that block RdRp (Sofia et al., 2012). For example, the cytidine analogue 4’-azidocytidine (R1479) and its prodrug balapiravir (R1626) have been assessed for treating HCV, and later dengue virus infections (Chen et al., 2014; Klumpp et al., 2006). However, balapiravir failed in clinical trials against dengue due to limited efficacy (Nguyen et al., 2013).

To test whether a combination therapy approach could improve the efficacy of R1479, we treated cells with R1479 in combination with GSK983 and CPU. As expected, R1479 showed dose-dependent inhibition of dengue virus replication with an EC_50_ ∼32 μM (Figure S6*A*). Inhibition of either *de novo* or salvage pyrimidine biosynthesis alone did not potentiate the antiviral activity of R1479 or cause additional cytotoxicity (Figure S6*A*). In contrast, inhibition of both the *de novo* and salvage pathways markedly enhanced the potency of R1479 (Figure 6*A*). In the presence of 250 μM CPU, the EC_50_ of R1479 was lowered to ∼12 μM. Notably, such a 3-component regimen had minimal impact on the cytotoxicity profile of R1479 (Figure 6*B*). Regarding other analogues, 5-F-CPU-GSK983 was able to slightly potentiate the antiviral activity of R1479 whereas 5’-F-CPU appeared to actually decrease antiviral efficacy of R1479 (Figure S6*B* and S6*C*).

**Figure 6.**
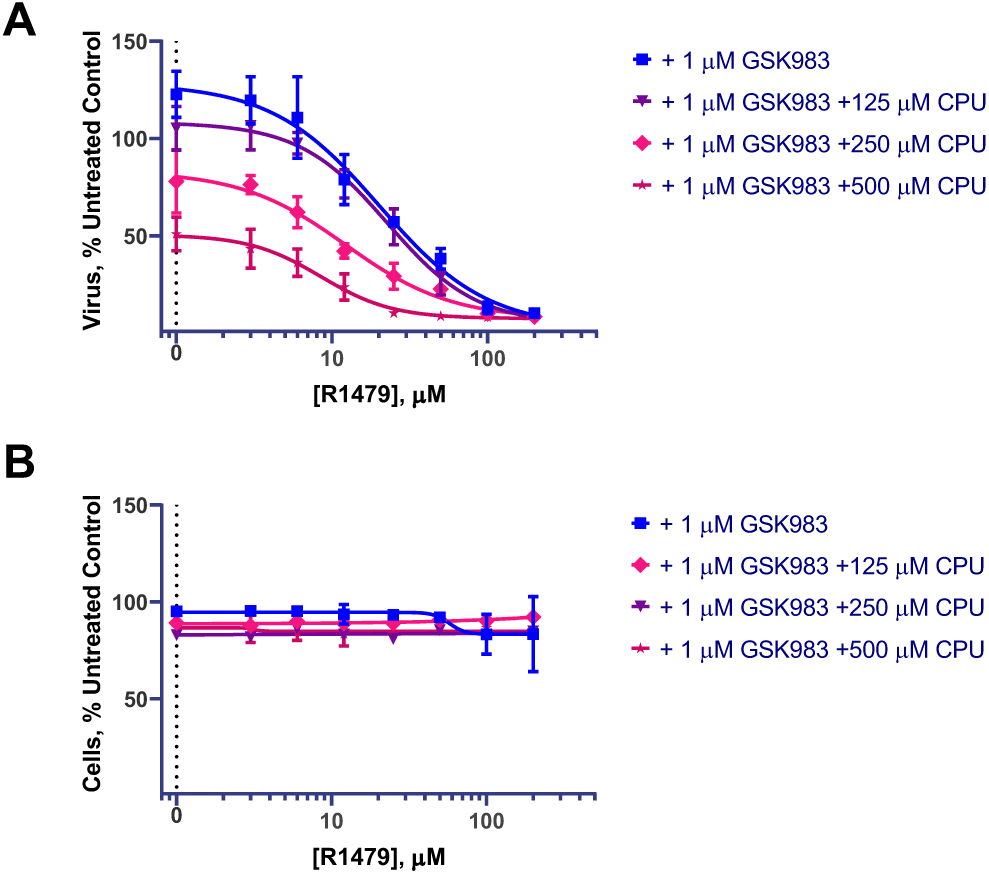
Potentiating the effects of RdRp inhibitor R1479 with CPU-GSK983 combination treatment. (*A*). Luminescence of DENV-2 infectious clone 16681 expressing luciferase in A549 cells at 48 h. (*B*). Cell viability of A549 cells under the triple combination therapy at 48 h. In all assays, the culture medium was supplemented with 20 μM uridine to mimic plasma uridine concentration and 1 μM GSK983 to block *de novo* pyrimidine biosynthesis. Error bars represent ± S.D. of three replicates.

As robust intracellular phosphorylation is critical for both R1479 and CPU activity, we confirmed that competition between these two compounds for intracellular kinases at the rate-limiting mono-phosphorylation step (Ford et al., 1991; Klumpp et al., 2008) should be minimal. Despite its 2’-α-OH group, R1479 is known to be solely phosphorylated by deoxycytidine kinase (dCK) (Klumpp et al., 2008) and was indeed not a UCK2 substrate in our hands (Figure S7). Similarly, cyclopentenyl compounds are substrates of UCK2 and not dCK (Ford et al., 1991).

To gain insight into the mechanism underlying the improvement of R1479 efficacy with CPU, we deployed a replicon assay that bypasses viral entry by electroporation of DENV RNA (Alvarez et al., 2005; Marceau et al., 2016). Such replicon systems include the coding region needed for viral RNA translation and replication, while lacking the structural genes necessary for the production of viral particles. Consequently, viral genome replication is examined independent of viral entry and particle assembly (Kato and Hishiki, 2016). In this assay (Figure 7), the EC_50_ of R1479 alone was ∼90 μM. Inhibition of *de novo* pyrimidine biosynthesis alone with GSK983 did not alter this value (∼96 μM), whereas addition of 250 μM CPU shifted it to ∼56 μM. Targeting both the *de novo* and salvage pathways with a combination of 1 μM GSK and 250 μM CPU lowered the EC_50_ of R1479 by over 4-fold to ∼19 μM. These results suggest that intracellular nucleotide depletion is the major driver of CPU’s antiviral activity. Together, these findings highlight the potential of combining modulators of pyrimidine metabolism with inhibitors of viral genome replication.

**Figure 7.**
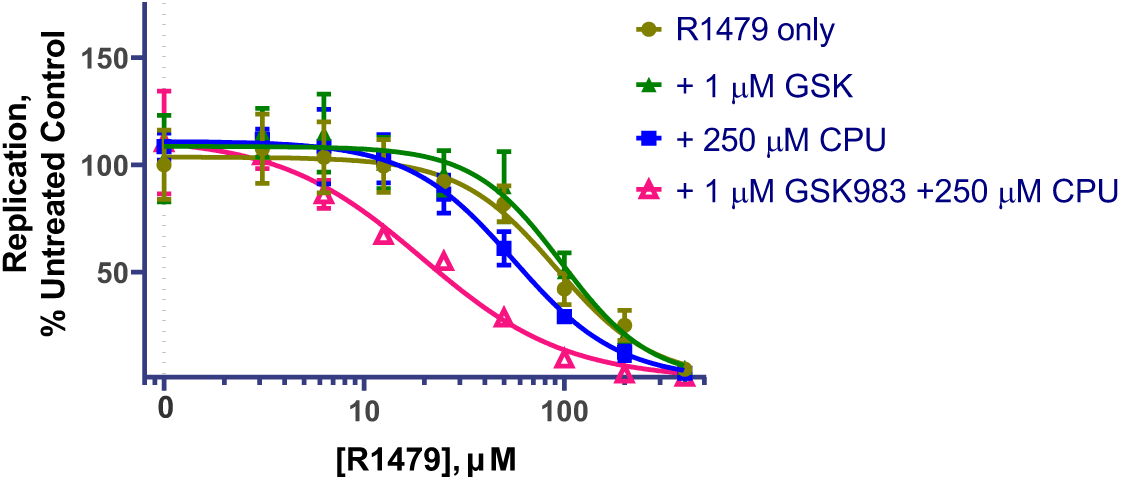
Examining viral replication with combinations of RdRp inhibitor R1479 with CPU-GSK983 treatment. (*A*). Luminescence of DENV replicon RNA expressing luciferase in A549 cells at 48 h. In all assays, the culture medium was supplemented with 20 μM uridine and 1 μM GSK983. Error bars represent ± S.D. of three replicates.

## DISCUSSION

The flaviviruses, including dengue virus, are an important class of clinically-relevant viral pathogens with limited treatment options (Boldescu et al., 2017). In particular, dengue causes hundreds of millions of symptomatic infections annually yet lacks any antiviral treatment while the sole approved vaccine has limited use because of safety risks associated with vaccination of individuals not previously exposed to dengue (Lim et al., 2013; Sridhar et al., 2018). A series of high-throughput phenotypic cell-based screens have recently identified DHODH inhibition as a potent, broad-spectrum antiviral strategy *in vitro* (Hoffmann et al., 2011; Lucas-Hourani et al., 2013; Luthra et al., 2018; Wang et al., 2011). Furthermore, a DHODH inhibitor was recently shown to suppress SARS-CoV-2 *in vitro*(Xiong et al., 2020). However, DHODH inhibitors often lose antiviral activity upon extracellular uridine addition due presumably to pyrimidine salvage (Deans et al., 2016).

While a prior study has explored combining a DHODH inhibitor with guanosine analogs ribavirin or INX-08189 in a non-uridine supplemented cell culture dengue model(Yeo et al., 2015), the antiviral efficacy of a DHODH and a salvage inhibitor has not been previously reported despite interest (Luthra et al., 2018). Herein, we found that a GSK983-CPU combination treatment effectively blocked dengue virus replication in the presence of physiological concentrations of extracellular uridine. At the same time, depletion of UTP and CTP led to a synergy between the CPU-GSK983 combination and RdRp inhibition.

Synthesis and evaluation of a series of CPU analogs revealed that CPU and 5-F-CPU were substrates of UCK2, and the resulting monophosphates were substrates of CMPK1. In combination with GSK983, the flux of CPU and 5-F-CPU to their triphosphate forms depleted intracellular pools of pyrimidine NTPs (Figure 4A). As the non-substrate analogue 5’-F-CPU did not deplete intracellular pyrimidines despite *in vitro* UCK2 inhibitory activity, our results suggest that CPU’s activity in host cells is driven by its ability to block multiple targets in pyrimidine salvage rather than solely UCK2 inhibition.

This distinctive antiviral activity of CPU could conceivably originate from both host-targeting modulation of pyrimidine metabolism as well as potential direct acting antiviral activity of its triphosphate on viral replication. Intracellular nucleotide depletion appears correlated with antiviral activity given that the structurally unrelated nucleoside transport inhibitor DPY also suppressed intracellular nucleotide levels and viral replication (Figure S5). Additionally, while 250 µM CPU in the absence of GSK983 did slightly potentiate the effects of R1479 on viral replication (Figure 7), this did not translate to decreased viral infection at the same time point (Figure S6*A*). Nevertheless, further investigation of inhibitory effects of CPU-TP on dengue RdRp is warranted. Indeed, as noted previously, NIs are the largest class of RdRp inhibitors (Klumpp et al., 2008). Additionally, a multi-factorial approach that incorporates information about interactions with multiple host and viral enzymes will likely be useful for future SAR around CPU as this compound remained the most efficacious in our panel.

CPU is known to be safe *in vitro* (Blaney et al., 1992; Ford et al., 1991; Song et al., 2001), and does not show appreciable toxicity in mice (Cysyk et al., 1995) or non-human primates (Blaney et al., 1990). While the safety profile of GSK983 remains less clear, other DHODH inhibitors like teriflunomide and brequinar have been extensively studied in humans, and are generally well-tolerated (Aly et al., 2017). Meanwhile, many RdRp inhibitors have been evaluated in clinical trials against dengue. While a preliminary study combining CPU with a *de novo* synthesis inhibitor *in vivo* resulted in toxicity (Cysyk et al., 1995), there is precedent for combining inhibitors of *de novo* pyrimidine synthesis with nucleoside transport inhibitors in humans(Casper et al., 1991). With careful dose optimization, our results suggest that a combination strategy targeting both host pyrimidine biosynthesis and viral RdRp holds promise as a potential therapy against RNA viruses. In light of the growing interest in RdRp and DHODH inhibitors as antiviral agents, our findings shine light on a promising way to enhance their clinical utility by combining them with modulators of mammalian pyrimidine metabolism.

## ACKNOWLEDGMENTS

The authors wish to thank Roberto Mateo, David Constant and Khanh Nguyen for helpful discussions and technical advice. This research was supported by a grant from the National Institutes of Health (1U19 AI109662 from the National Institute of Allergy and Infectious Diseases) to C.K., M.B., and J.C. Also, J.C. acknowledges NIH (R01 AI141970) for financial support. A.G. is supported by a National Science Foundation Graduate Research Fellowship and a Stanford ChEM-H Chemistry/Biology Interface Predoctoral Training Program Fellowship.

## AUTHOR CONTRIBUTIONS

Q.L, A.G. and C.K. designed research; Q.L., A.G. and W.Q. performed research; Q.L, A.G., A.O., C.F., M.S., J.C., M.B. and C.K. analyzed data; Q.L., A.G. and C.K. wrote the paper.

## MATERIALS AND METHODS

### CELL CULTURE

All cell lines were maintained in a humidified incubator (37°C, 5% CO_2_). A549 cells were obtained from ATCC and cultured in DMEM supplemented with 10% FBS, penicillin/streptomycin, and L-glutamine. Upon reaching 50-75% confluence, A549 cells were detached from the growth surface using a trypsin/EDTA solution prior to analysis. Cells were maintained in logarithmic growth during all biological assays.

### DENV-LUC REPORTER VIRUS GENERATION

The design of the pDENV-Luc infectious clone derived from dengue serotype 2 (DENV-2) strain 16681 was described in detail by Marceau *et al* (Marceau et al., 2016). The plasmid was linearized with XbaI and in vitro transcribed into the genomic RNA of DENV-Luc virus using the T7 Megascript Kit (Ambion) in the presence of m7G(5’)ppp(5’)G RNA Cap Structure Analog (NEB). 5 μg DENV-Luc RNA was electroporated into 2×10^6^ Vero cells. The transfected cells were resuspended with Dulbecco’s modified Eagle’s medium (DMEM; Invitrogen) with 10% FBS and 100 U/ml penicillin-streptomycin and transferred into a T-175 flask incubated at 37 °C with 5% CO_2_. Supernatants were collected and replenished with fresh medium every 24 h from day 17 to 24 post-transfection, pooled together, clarified by centrifugation and stored in aliquots at −80°C. The amount of infectious DENV-Luc virus in the stock was titrated using the TCID_50_ assay and calculated by the Spearman & Kärber algorithm as described previously (Hierholzer and Killington, 1996).

### pDENV-LUC REPLICON GENERATION

The plasmid pDENV-Luc replicon containing a Renilla luciferase expressing DENV subgenomic replicon was described previously (Marceau et al., 2016). For *in vitro* transcription, 10 ug of plasmid DNA was linearized using XbaI restriction enzyme. Replicon RNA was generated using the MEGAscript T7 Transcription Kit in the presence of 5mM m7G(5′)ppp(5′)G RNA Cap Structure Analogue. Two million cells were washed twice with PBS and re-suspended in electroporation buffer (Teknova, E0399), mixed with 4 ug of purified replicon RNA, and electroporated using Bio-Rad Gene Pulser Xcell electroporator using square wave protocol. Electroporated cells were resuspended in pre-warmed culture medium without antibiotics and seeded as required. Luciferase expression was measured using Renilla Luciferase Assay system (Promega, E2820) following the manufacturers’ instructions.

### GENERAL MATERIALS AND METHODS FOR CHEMICAL SYNTHESES

Unless otherwise noted, all reactions were performed under an argon atmosphere in flame- or oven-dried glassware. Reaction mixtures were stirred using Teflon-coated magnetic stirrer bars and monitored by thin layer silica gel chromatography (TLC) using 0.25 mm silica gel 60F plates with fluorescent indicator from Merck. Plates were visualized under UV or treated by KMnO4 stain with gentle heating. Products were purified on an AnaLogix IntelliFlash 280 Flash column chromatography system using the solvent gradients indicated. Anhydrous tetrahydrofuran (THF), dichloromethane (CH_2_Cl_2_), dimethylformamide (DMF), acetone, and dimethyl sufoxide (DMSO) were obtained from Acros Organics. Diethyl ether (Et_2_O), ethyl acetate (EtOAc), hexanes, and methanol (MeOH) were from Fisher Scientific. DMSO used in bioassays and to prepare biological samples was from Fisher BioReagents. All other reagents were from commercial suppliers and were used as received without additional purification. Samples prepared for biological evaluation were purified via preparative HPLC in a water/acetonitrile (MeCN) gradient containing 0.1% (v/v) trifluoracetic acid (TFA) using an Agilent 1260 Infinity system equipped with an Agilent Prep-C18 column (21.2 × 250 mm).

NMR spectra were measured on a Varian INOVA 500 (^1^ H at 500 MHz, ^13^C at 125 MHz), a Varian 400 (^1^H at 400 MHz, ^13^C at 100 MHz), or a Varian INOVA 600 MHz (^1^H at 500 MHz, ^13^C at 150 MHz) magnetic resonance spectrometer, as noted. ^1^H chemical shifts were reported relative to the residual solvent peak (CDCl_3_= 7.26 ppm; MeOD = 3.31 ppm) as follows: chemical shift (δ) [multiplicity (s = singlet, brs=broad singlet, d = doublet, t = triplet, q = quartet, dd = doublet of doublet, m = multiplet), coupling constant(s) in Hz, integration]. ^13^C chemical shifts were reported relative to the residual deuterated solvent ^13^C signals (CDCl_3_ = 77.16 ppm, MeOD = 49.00 ppm) and rounded to one decimal places. Infrared spectra were recorded on a Nicolet iS50 FT/IR Spectrometer at the Stanford Nano Share Facilities and were reported in wavenumbers (cm^−1^). Optical rotation data were obtained using a JASCO DIP were reported as 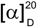(c = grams/100 mL, solvent), where D indicates the sodium D line (589 nm). High resolution mass spectra were obtained on an Agilent 6545 QTof mass spectrometer at the Metabolic Chemistry Analysis Center at Stanford University.

## METHOD DETAILS

### Synthesis of CPU analogs and compound characterization

#### 3-benzoyl-1-((3a*S*,4*R*,6a*R*)-6-(((*tert*-butyldiphenylsilyl)oxy)methyl)-2,2-dimethyl-3a,6a-dihydro-4*H*-cyclopenta[*d*][1,3]dioxol-4-yl)pyrimidine-2,4(1*H*,3*H*)-dione (3a)

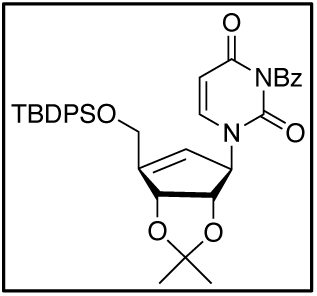

Compound **1** was prepared in 10 steps from D-ribose according to literature procedures (Choi et al., 2004). To a suspension of **1** (440 mg, 1.04 mmol, 1 equiv.), **2a** (270 mg, 1.25 mmol, 1.2 equiv.), and PPh_3_ (410 mg, 1.56 mmol, 1.5 equiv.) in THF (12 mL) at 0 °C was added DEAD solution in toluene (720 μL, 1.5 equiv., 40 w% in toluene). After the addition of DEAD, the reaction mixture turned from a white suspension to a yellow solution, which was slowly warmed up to room temperature and stirred overnight. The reaction mixture was then concentrated, loaded onto a 12g SiO_2_ flash cartridge, and purified with a linear gradient of 20-40% EtOAc in hexanes to afford **3a** (446 mg, 0.72 mmol, 69%) as white powder. 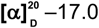 (*c* 1.3, MeOH); **IR** (film, cm^−1^) ;3071, 2930, 2857, 1748, 1705, 1667, 1441, 1429, 1372, 1234, 1112, 9055, 732, 704.^**1**^**H NMR** (400 MHz, CDCl_3_) δ 7.95 (d, *J* = 7.4 Hz, 1H), 7.71 – 7.61 (m, 6H), 7.53 – 7.47 (m, 2H), 7.45 – 7.42 (m, 2H), 7.41 – 7.34 (m, 4H), 6.91 (d, *J* = 8.1 Hz, 1H), 5.77 (d, *J* = 8.1 Hz, 1H), 5.67 (s, 1H), 5.37 (s, 1H), 5.08 (d, *J* = 5.8 Hz, 1H), 4.60 (d, *J* = 5.8 Hz, 1H), 4.47 (d, *J* = 16.4 Hz, 1H), 4.41 (d, *J* = 16.7 Hz, 1H), 1.34 (s, 3H), 1.29 (s, 3H), 1.10 (s, 9H). ^**13**^**C NMR** (100 MHz, CDCl_3_) δ 168.8, 162.3, 153.6, 149.8, 141.1, 135.6, 135.6, 135.3, 133.3, 133.1, 131.6, 130.7, 130.1, 129.3, 128.0, 128.0, 120.8, 112.9, 102.4, 84.6, 83.4, 68.4, 61.4, 27.3, 27.0, 25.9, 19.4. **HRMS** (ESI) *m/z* 623.2574 [(M+H)^+^; calcd for C_36_H_39_N_2_O_6_Si^+^: 623.2572].

#### 1-((1*R*,4*R*,5*S*)-4,5-dihydroxy-3-(hydroxym ethyl)cyclopent-2-en-1-yl)pyrimidine-2,4(1*H*,3*H*)-dione (4a)

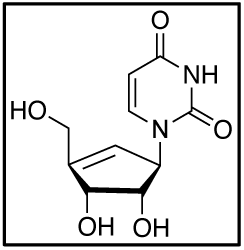

To compound **3a** (30 mg, 0.048 mmol) was added 0.5 mL 7N NH_3_ solution in methanol. After 1 h, the reaction solvent was removed under positive N_2_ atmosphere and the resulting solid was further dried under high vacuum. The reaction crude was then treated with 30 μL HCl in 300 μL THF and stirred overnight. Excess NaHCO_3_ was added to neutralize the reaction, followed by the addition of 1 mL MeOH. The resulting suspension was filtered through a short Celite pad, rinsed with MeOH, and concentrated. The resulting crude was then resuspended with 1 mL H_2_O and subjected to HPLC purification with a linear gradient of 5-20% MeCN (0.1% TFA) in H_2_O (0.1% TFA) on a prep C18 column (Agilent 10 prep-C18 250 × 21.1 mm). Fractions containing desired products was then combined and lyophilized to afford the final product **4a** (4.6 mg, 0.019 mmol, 40%) as white powder. 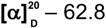 (*c* 3.2, MeOH); **IR** (film, cm^−1^): 3349 (br), 1667, 1465, 1390, 1258, 1202, 1114.^**1**^**H NMR** (500 MHz, MeOD) δ 7.42 (d, *J* = 7.9 Hz, 1H), 5.71 (s, 1H), 5.49 (s, 1H), 4.90 (s, 1H), 4.52 (d, *J* = 5.9, 1.2 Hz, 1H), 4.33 – 4.19 (m, 2H), 4.04 (t, *J* = 5.6 Hz, 1H). ^**13**^**C NMR** (125 MHz, MeOD) δ 166.4, 153.1, 152.1, 143.6, 125.5, 102.8, 78.4, 74.0, 67.4, 60.3**. HRMS** (ESI) *m/z* 263.0644 [(M+Na)^+^; calcd for C_10_H_12_N_2_NaO_5_^+^: 263.0638].

In a similar manner, compound 1 (360 mg, 0.85 mmol, 1 equiv.) was reacted with 2b (238 mg, 1.02 mmol, 1.2 equiv.) to afford intermediate 3b as white powder (288 mg, 0.45 mmol, 53%). Intermediate **3b** (32 mg, 0.05 mmol) was then deprotected to afford **4b** (7.7 mg, 0.03 mmol, 60%) as white powder.

#### 3-benzoyl-1-((3a*S*,4*R*,6a*R*)-6-(((*tert*-butyldiphenylsilyl)oxy)methyl)-2,2-dimethyl-3a,6a-dihydro-4*H*-cyclopenta[*d*][1,3]dioxol-4-yl)-5-fluoropyrimidine-2,4(1*H*,3*H*)-dione (3b)

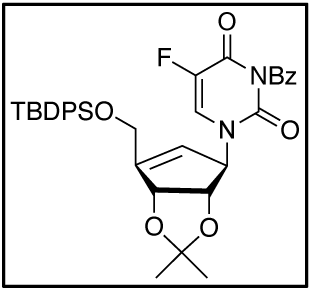

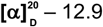 (*c* 1.4, CHCl_3_); **IR** (film, cm^−1^): 3072, 2932, 2857, 1753, 1712, 1665, 1662, 1448, 1373, 1234, 1106, 1088, 732, 702, 687. ^**1**^**H NMR** (400 MHz, CDCl_3_) δ 7.97 – 7.92 (m, 2H), 7.73 – 7.63 (m, 5H), 7.55 – 7.49 (m, 2H), 7.48 – 7.42 (m, 2H), 7.42 – 7.35 (m, 4H), 7.06 (d, *J* = 5.8 Hz, 1H), 5.68 (brs, 1H), 5.41 (brs, 1H), 5.08 (d, *J* = 5.8 Hz, 1H), 4.59 (dd, *J* = 12.1, 6.4 Hz, 1H), 4.54 – 4.35 (m, 2H), 1.34 (s, 3H), 1.29 (s, 3H), 1.11 (s, 9H). ^**13**^**C NMR** (100 MHz, CDCl_3_) δ 167.3, 154.3, 148.3, 141.4, 139.0, 135.6, 135.6, 133.2, 133.0, 131.2, 130.8, 130.2, 130.2, 129.4, 129.1, 128.0, 125.5, 125.1, 120.4, 113.0, 84.6, 83.2, 68.3, 61.4, 27.3, 27.0, 25.9, 19.4. ^19^F NMR (376 MHz, CDCl_3_) δ −163.44 (d, *J* = 5.7 Hz). **HRMS** (ESI) *m/z* 663.2323 [(M+Na)^+^; calcd for C_36_H_37_FN_2_O_6_SiNa^+^:663.2297].

#### 1-((1*R*,4*R*,5*S*)-4,5-dihydroxy-3-(hydroxymethyl)cyclopent-2-en-1-yl)-5-fluoropyrimidine-2,4(1*H*,3*H*)-dione (4b)

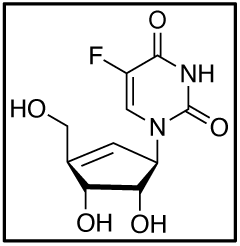

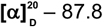 (*c* 0.45, MeOH); **IR** (film, cm^−1^): 3375 (br), 1696, 1660, 1386, 1242, 1115, 1011.^**1**^**H NMR** (400 MHz, MeOD) δ 7.60 (d, *J* = 6.6 Hz, 1H), 5.68 (q, *J* = 1.8 Hz, 1H), 5.49 (brs, 1H), 4.52 (d, *J* = 5.6 Hz, 1H), 4.26 (d, *J* = 2.2 Hz, 2H), 4.03 (t, *J* = 5.5 Hz, 1H). ^**13**^**C NMR** (100 MHz, MeOD) δ 159.54 (d, *J* = 26.1 Hz), 152.64, 151.74, 142.07 (d, *J* = 233.0 Hz), 127.39 (d, *J* = 33.7 Hz), 125.24, 78.17, 73.94, 67.73, 60.26. ^**19**^**F NMR** (376 MHz, MeOD) δ −168.53 (dd, *J* = 6.8, 1.7 Hz).**HRMS** (ESI) *m/z* 259.0722 [(M+H)^+^; calcd for C_10_H_12_FN_2_O_5_ ^+^: 259.0725].

In a similar manner, compound **1** (245 mg, 0.58 mmol, 1 equiv.) was reacted with **2c** (238 mg,0.70 mmol, 1.2 equiv.) to afford intermediate **3c** (160 mg, 0.21mmol, 36%) as white powder. Intermediate **3c** (40 mg, 0.053 mmol) was then deprotected to afford **4c** (9.6 mg, 0.026 mmol, 49%) as white powder.

#### 3-benzoyl-1-((3a*S*,4*R*,6a*R*)-6-(((*tert*-butyldiphenylsilyl)oxy)methyl)-2,2-dimethyl-3a,6a-dihydro-4*H*-cyclopenta[*d*][1,3]dioxol-4-yl)-5-iodopyrimidine-2,4(1*H*,3*H*)-dione (3c)

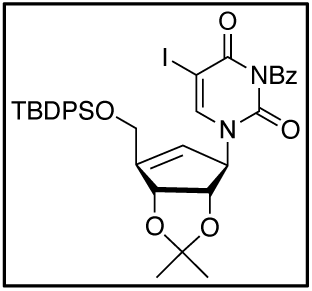

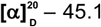 (*c* 1.3, CHCl_3_); **IR** (film, cm^−1^): 3071, 2932, 2857, 1749, 1703, 1666, 1608, 1420, 1233, 1112, 703.^**1**^**H NMR** (400 MHz, CDCl_3_) δ 7.93 – 7.90 (m, 1H), 7.72 – 7.63 (m, 5H), 7.54 – 7.48 (m, 3H), 7.45 – 7.37 (m, 6H), 5.72 (brs, 1H), 5.37 (s, 1H), 5.13 (d, *J* = 5.8 Hz, 1H), 4.62 (d, *J* = 5.8 Hz, 1H), 4.51 – 4.34 (m, 2H), 1.34 (s, 3H), 1.29 (s, 3H), 1.09 (s, 9H). ^**13**^**C NMR** (100 MHz, CDCl_3_) δ 167.8, 159.1, 154.1, 149.5, 145.8, 135.6, 135.4, 133.1, 133.1, 131.1, 130.7, 130.1, 129.4, 128.0, 128.0, 120.6, 113.0, 84.6, 83.4, 69.2, 68.1, 61.4, 27.3, 27.0, 25.9, 19.5. **HRMS** (ESI) *m/z* 749.1538 [(M+H)^+^; calcd for C_36_H_38_IN_2_O_6_Si^+^:749.1538].

#### 1-((1*R*,4*R*,5*S*)-4,5-dihydroxy-3-(hydroxymethyl)cyclopent-2-en-1-yl)-5-iodopyrimidine-2,4(1*H*,3*H*)-dione (4c)

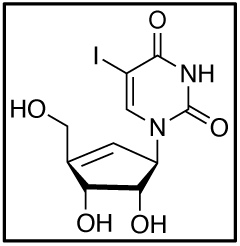

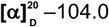 (*c* 0.6, MeOH); **IR** (film, cm^−1^): 3370 (br), 1682, 1608, 1424, 1260, 1111.^**1**^**H NMR** (400 MHz, MeOD) δ 7.78 (s, 1H), 5.70 (q, *J* = 1.8 Hz, 1H), 5.47 (brs, 1H), 4.52 (d, *J* = 5.5 Hz, 1H), 4.33-4.20 (m, *J* = 2.1 Hz, 2H), 4.04 (t, *J* = 5.7 Hz, 1H).^**13**^**C NMR** (100 MHz, MeOD) δ 163.0, 152.8, 152.4, 148.0, 125.5, 78.6, 73.9, 68.6, 67.9, 60.0.**HRMS** (ESI) *m/z* 366.9782 [(M+H)^+^; calcd for C_10_H_12_IN_2_O_5_ ^+^: 366.9785].

In a similar manner, compound **1** (113 mg, 0.27 mmol, 1 equiv.) was reacted with **2d** (74 mg, 0.32 mmol, 1.2 equiv.) to afford intermediate **3d** as white powder (100 mg, 0.16 mmol, 59%). Intermediate **3d** (30 mg, 0.047 mmol) was then deprotected to afford **4d** (4.3 mg, 0.017 mmol, 36%) as white powder.

#### 3-benzoyl-1-((3a*S*,4*R*,6a*R*)-6-(((*tert*-butyldiphenylsilyl)oxy)methyl)-2,2-dimethyl-3a,6a-dihydro-4*H*-cyclopenta[*d*][1,3]dioxol-4-yl)-5-methylpyrimidine-2,4(1*H*,3*H*)-dione (3d)

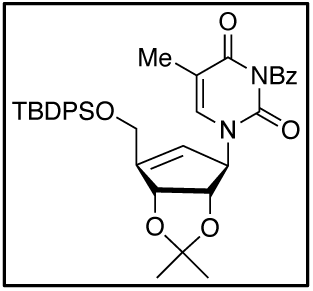

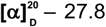 (*c* 1.2, CHCl_3_); **IR** (film, cm^−1^): 3071, 2932, 2857, 1748, 1699, 1656, 1429, 1371, 1235, 1111, 732, 702.^**1**^**H NMR** (500 MHz, CDCl3) δ 7.97 – 7.89 (m, 2H), 7.71 – 7.61 (m, 5H), 7.53 – 7.46 (m, 3H), 7.45 – 7.42 (m, 1H), 7.40 – 7.35 (m, 4H), 6.90 (brs, 1H), 5.73 (s, 1H), 5.37 (s, 1H), 5.13 (d, *J* = 5.8 Hz, 1H), 4.65 (dd, *J* = 16.3, 4.7 Hz, 1H), 4.47 – 4.37 (m, 2H), 1.96 (s, 3H), 1.34 (s, 3H), 1.29 (s, 3H), 1.09 (s, 9H).^**13**^**C NMR** (125 MHz, CDCl_3_) δ 169.1, 163.0, 152.9, 149.8, 137.2, 135.6, 135.1, 133.2, 133.1, 131.7, 130.6, 130.1, 129.3, 128.0, 121.3, 112.7, 111.0, 84.6, 83.5, 68.3, 61.4, 27.3, 27.0, 25.9, 19.5, 12.8. **HRMS** (ESI) *m/z* 637.2734 [(M+H)^+^; calcd for C_37_H_41_N_2_O_6_Si^+^: 637.2728].

#### 1-((1*R*,4*R*,5*S*)-4,5-dihydroxy-3-(hydroxymethyl)cyclopent-2-en-1-yl)-5-methylpyrimidine-2,4(1*H*,3*H*)-dione (4d)

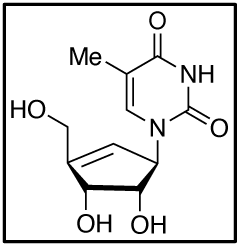

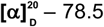 (*c* 0.38, MeOH); **IR** (film, cm^−1^): 3371, 1683, 1476, 1261, 1205, 1114, 1016.^**1**^**H NMR** (400 MHz, MeOD) δ 7.23 (d, *J* = 1.2 Hz, 1H), 5.68 (q, *J* = 1.8 Hz, 1H), 5.48 (brs, 1H), 4.52 (d, *J* = 5.9 Hz, 1H), 4.26 (q, *J* = 2.5 Hz, 2H), 4.04 (t, *J* = 5.6 Hz, 1H), 1.87 (d, *J* = 1.2 Hz, 3H). ^**13**^**C NMR** (100 MHz, MeOD) δ 166.5, 153.3, 151.8, 139.2, 126.0, 111.8, 78.3, 74.0, 67.2, 60.3, 12.3. **HRMS** (ESI) *m/z* 277.0799 [(M+Na)^+^; calcd for C_11_H_14_N_2_O_5_Na^+^: 277.0795].

#### 3-benzoyl-1-((3a*S*,4*R*,6a*R*)-6-(hydroxymethyl)-2,2-dimethyl-3a,6a-dihydro-4*H*-cyclopenta[*d*][1,3]dioxol-4-yl)pyrimidine-2,4(1*H*,3*H*)-dione (5)

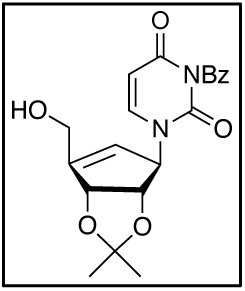

To a solution of **3a** (300 mg, 0.48 mmol, 1 equiv.) in THF (5 mL) was added TBAF (580 μL, 0.58 mmol, 1.2 equiv., 1 M in THF) at 0 °C. After 2 h, the reaction mixture was quenched with 5 mL of saturated NH_4_OH solution and extracted with 5 5 mL EtOAc. The combined organic layers were dried over anhydrous Na_2_SO_4_, concentrated, loaded onto a 4 g SiO_2_ flash cartridge, and purified with a linear gradient 80-95% EtOAc in hexanes to afford the free primary alcohol **5** (160 mg, 0.42 mmol, 87%) as white powder. 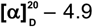 (*c* 4.5, CHCl_3_); **IR** (film, cm^−1^): 3473 (br), 3088, 2988, 2933, 1746, 1702, 1665, 1443, 1374, 1239, 1179, 1239, 1059.^**1**^**H NMR** (500 MHz, CDCl_3_) δ 7.93 (d, *J* = 8.4 Hz, 2H), 7.70 – 7.61 (m, 1H), 7.50 (dd, *J* = 8.2, 7.4 Hz, 2H), 7.16 (d, *J* = 8.0 Hz, 1H), 5.81 (d, *J* = 8.0 Hz, 1H), 5.64 (s, 1H), 5.30 (s, 1H), 5.25 (d, *J* = 5.8 Hz, 1H), 4.70 (d, *J* = 5.8 Hz, 1H), 4.43 (d, *J* = 15.9 Hz, 1H), 4.38 (d, *J* = 15.9 Hz, 1H), 1.43 (s, 3H), 1.34 (s, 3H). ^**13**^**C NMR** (125 MHz, CDCl_3_) δ 168.8, 162.2, 152.5, 149.7, 141.6, 135.3, 131.5, 130.7, 129.3, 121.7, 113.0, 102.5, 84.3, 84.0, 69.2, 60.2, 27.3, 25.8. **HRMS** (ESI) *m/z* 385.1391 [(M+H), calcd for C_20_H_21_N_2_O_6_ ^+^: 385.1394].

#### 1-((1*R*,4*R*,5*S*)-4,5-dihydroxy-3-(methoxymethyl)cyclopent-2-en-1-yl)pyrimidine-2,4(1*H*,3*H*)-dione (6)

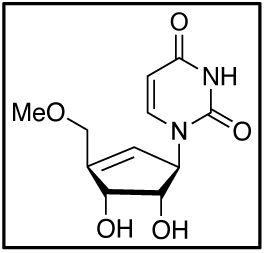

To a solution of **5** (20 mg, 0.05 mmol, 1 equiv.) in dry acetone (0.5 mL) was added Ag_2_O (23 mg, 0.1 mmol, 2 equiv.) and MeI (31 μL, 0.5 mmol, 10 equiv.). The reaction mixture was stirred at room temperature for 24 h, filtered through Celite pad, and concentrated in vacuo to afford a residue oil 17 mg. In a similar manner as the synthesis of **3a** from **4a**, the crude was treated with 0.5 mL 7N NH_3_ in methanol, followed by 25 μL HCl in 250 THF to furnish the analog **6** (2.5 mg, 0.01 mmol, 20% for three steps). 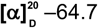 7 (*c* 0.25, MeOH); **IR** (film, cm^−1^) ; 3369 (br), 2921, 2851, 1682, 1469, 1410, 1262, 1204, 1096.^**1**^**H NMR** (600 MHz, MeOD) δ 7.40 (d, *J* = 8.0 Hz, 1H), 5.72 (dd, *J* = 3.8, 1.7 Hz, 1H), 5.69 (d, *J* = 8.0 Hz, 1H), 5.51 – 5.43 (m, 1H), 4.50 (d, *J* = 5.8 Hz, 1H), 4.14 (ddd, *J* = 14.1, 2.5, 2.4 Hz, 1H), 4.08 (ddd, *J* = 14.2, 2.2, 1.8 Hz, 1H), 4.05 (dd, *J* = 5.7, 5.5 Hz, 1H), 3.40 (s, 3H). ^**13**^**C NMR** (100 MHz, MeOD) δ 166.4, 153.0, 148.8, 143.7, 127.5, 102.8, 78.1, 74.0, 70.3, 67.6, 59.0. **HRMS** (ESI) *m/z* 277.0799 [(M+Na)^+^; calcd for C_11_H_14_N_2_O_5_Na^+^: 277.0795].

#### 3-benzoyl-1-((3a*S*,4*R*,6a*R*)-6-(fluoromethyl)-2,2-dimethyl-3a,6a-dihydro-4*H*-cyclopenta[*d*][1,3]dioxol-4-yl)pyrimidine-2,4(1*H*,3*H*)-dione (S1)

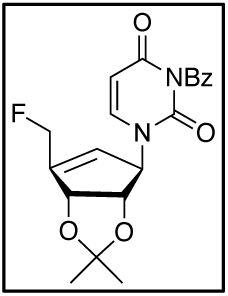

To alcohol **5** (50 mg, 0.13 mmol, 1 equiv.) in 1 mL CH_2_Cl_2_ at –78°C was added a stock solution of DAST (0.15 mmol, 1.2 equiv., 200 μL, prepared by dilute 100 μL of DAST with CH_2_Cl_2_ to 1 mL). The reaction mixture was warmed up to room temperature and stirred overnight before quenching with 1 mL saturated NaHCO_3_. The resulting mixture was then extracted with 3×3 mL EtOAc. The combined organic layers were dried over anhydrous Na_2_SO_4_, concentrated, loaded onto a 4 g SiO_2_ flash cartridge, and purified with a linear gradient 20-50% EtOAc in hexanes to afford the intermediate **S1** (27 mg, 0.070 mmol, 54%) as white powder. 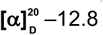 (*c* 2.4, CHCl_3_); **IR** (film, cm^−1^): 3100, 2989, 2937, 1746, 1704, 1667, 1599, 1441, 1374, 1239, 1179, 1060,989.^**1**^**H NMR** (600 MHz, CDCl3) δ 7.93 (d, *J* = 8.1 Hz, 2H), 7.66 (t, *J* = 7.5 Hz, 1H), 7.51 (dd, *J* = 7.91, 7.13 Hz, 2H), 7.13 (d, *J* = 8.1 Hz, 1H), 5.83 (d, *J* = 8.1 Hz, 1H), 5.72 (s, 1H), 5.32 (s, 1H), 5.26 (d, *J* = 5.9 Hz, 1H), 5.14 (s, 1H), 5.07 (s, 1H), 4.73 (d, *J* = 5.7 Hz, 1H), 1.43 (s, 3H), 1.34 (s, 3H). ^**13**^**C NMR** (125 MHz, CDCl_3_) δ 168.7, 162.2, 149.6, 141.5, 135.4, 131.5, 130.7, 129.4, 123.2, 113.2, 102.7, 95.7, 84.1, 83.2, 80.5 (d, *J* = 149 Hz), 78.8, 77.4, 77.2, 76.9, 69.2, 27.3, 25.8. ^**19**^**F NMR** (376 MHz, CDCl_3_) δ – 224.4 (t, *J* = 46.5). **HRMS** (ESI) *m/z* 387.1366 [(M+H)^+^; calcd for C_20_H_20_FN_2_O_5_^+^: 387.1351].

#### 1-((1*R*,4*R*,5*S*)-3-(fluoromethyl)-4,5-dihydroxycyclopent-2-en-1-yl)pyrimidine-2,4(1*H*,3*H*)-dione (7)

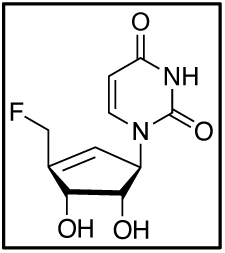

In a similar manner as the synthesis of **4a** from **3a**, compound **S1** (13 mg, 0.034 mmol, 1 equiv.) was treated with 0.5 mL 7N NH_3_ in methanol, followed by 25 μL HCl in 250 μL THF to furnish the analog **7** (4.4 mg, 0.018 mmol, 53% for two steps) as white powder. 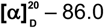 (*c* 0.22, MeOH); **IR** (film, cm^−1^): 3365 (br), 2920, 2852, 1675, 1466, 1389, 1265, 1203, 1115.^**1**^**H NMR** (600 MHz, MeOD) δ 7.42 (d, *J* = 7.9 Hz, 1H), 5.85 (brs, 1H), 5.70 (d, *J* = 8.0 Hz, 1H), 5.48 (brs, 1H), 5.15 – 4.99 (m, 2H), 4.57 (d, *J* = 6.0 Hz, 1H), 4.10 (dd, *J* = 6.7, 6.0 Hz, 1H). ^**13**^**C NMR** (150 MHz, MeOD) δ 166.2, 152.8, 147.2, 143.6, 128.2, 102.8, 80.5 (d, *J* = 149 Hz), 77.6, 73.1, 67.4. ^**19**^**F NMR** (376 MHz, MeOD) δ – 224.9 (t, *J* = 46.8). **HRMS** (ESI) *m/z* 265.0592 [(M+Na)^+^; calcd for C_10_H_11_FN_2_O_4_Na^+^: 265.0595].

#### ((3a*S*,4*R*,6a*R*)-4-(3-benzoyl-2,4-dioxo-3,4-dihydropyrimidin-1(2*H*)-yl)-2,2-dimethyl-3a,6a-dihydro-4*H*-cyclopenta[*d*][1,3]dioxol-6-yl)methyl methanesulfonate (8)

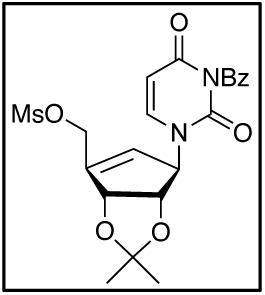

To a solution of **5** (60 mg,0.16 mmol, 1 equiv.) in anhydrous CH_2_Cl_2_ (2 mL) was added Et_3_N (44 μL, 0.32 mmol, 2 equiv.) at 0 °C, followed by the dropwise addition of 0.2 mL MsCl stock solution (0.26 mmol, 1.6 equiv.; Stock solution was prepared by diluting 0.1 mL MsCl with CH_2_Cl_2_ to 1 mL). After 20 min at 0 °C, the reaction mixture was quenched with 2 mL saturated aqueous solution of NH_4_Cl, extracted with 3 × 3 mL CH_2_Cl_2_, dried over anhydrous Na_2_SO_4_, concentrated, loaded on a 4g SiO_2_ flash cartridge, and purified with a linear gradient 50-95% EtOAc in hexanes to afford the compound **8** (53 mg, 0.11 mmol, 72%) as pale yellow oil. 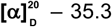 (*c* 2.8, CHCl_3_); **IR** (film, cm^−1^): 2989, 2937, 1745, 1702, 1661, 1597, 1440, 1354, 1235, 1174, 1087, 959, 906, 729.^**1**^**H NMR** (500 MHz, CDCl_3_) δ 7.92 (d, *J* = 7.2 Hz, 2H), 7.66 (dd, *J* = 8.0, 7.7 Hz, 1H), 7.50 (d, *J* = 7.8 Hz, 2H), 7.19 (d, *J* = 8.1 Hz, 1H), 5.83 – 5.67 (m, 2H), 5.30 (d, *J* = 4.8 Hz, 2H), 4.98 (d, *J* = 14.0 Hz, 1H), 4.83 (d, *J* = 14.0 Hz, 1H), 4.71 (d, *J* = 5.8 Hz, 1H), 3.04 (s, 3H), 1.41 (s, 3H), 1.33 (s, 3H). ^**13**^**C NMR** (125 MHz, CDCl_3_) δ 168.7, 162.1, 149.6, 146.3, 141.8, 135.4, 131.4, 130.6, 129.4, 125.9, 113.1, 102.7, 83.8, 83.1, 77.4, 77.2, 76.9, 68.7, 65.0, 38.1, 27.3, 25.7. **HRMS** (ESI) *m/z* 463.1185 [(M+H)^+^; calcd for C_21_H_23_N_2_O_8_S^+^: 463.1170].

#### 1-((3a*S*,4*R*,6a*R*)-6-(azidomethyl)-2,2-dimethyl-3a,6a-dihydro-4*H*-cyclopenta[*d*][1,3]dioxol-4-yl)pyrimidine-2,4(1*H*,3*H*)-dione (S2)

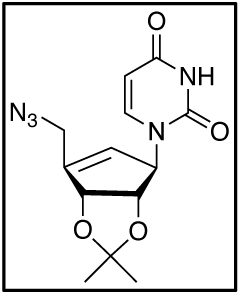

To the mesylate intermediate **8** (26 mg, 0.056 mmol, 1 equiv.) was added DMF (3 mL) and NaN_3_ (80 mg, 1.23 mmol, 22 equiv.). The resulting suspension was heated to 100°C and stirred for 18 hours, followed by the addition of 3 mL saturated solution of NH_4_Cl. The organic layer was extracted with 3 × 5 Et_2_O, washed with brine, and concentrated under reduced pressure. The crude material was purified through a 4g SiO_2_ flash cartridge with a linear gradient 40-95% EtOAc in hexanes to afford the azide intermediate **S2** (14 mg, 0.046 mmol, 82%) as brown oil. 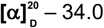 (*c* 0.7, CHCl_3_); **IR** (film,cm^−1^): 3197, 3059, 2988, 2929, 2102, 1687, 1456, 1380, 1243, 1082, 1058.^**1**^**H NMR** (500 MHz, CDCl_3_) δ 8.78 (brs, 1H), 7.02 (d, *J* = 8.0 Hz, 1H), 5.71 (d, *J* = 8.0 Hz, 1H), 5.64 (brs, 1H), 5.31 (brs, 1H), 5.22 (d, *J* = 5.7 Hz, 1H), 4.65 (d, *J* = 5.8 Hz, 1H), 4.13 (d, *J* = 16.0 Hz, 1H), 4.04 (d, *J* = 16.0 Hz, 1H), 1.44 (s, 3H), 1.35 (s, 3H). ^**13**^**C NMR** (125 MHz, CDCl_3_) δ 162.8, 150.5, 150.1, 141.2, 122.1, 112.6, 102.3, 86.2, 84.4, 68.0, 27.4, 26.0, 14.6. **HRMS** (ESI) *m/z* 306.1195 [(M+H)^+^; calcd for C_13_H_16_N_5_O_4_ ^+^: 306.1197].

#### 1-((1*R*,4*R*,5*S*)-3-(azidomethyl)-4,5-dihydroxycyclopent-2-en-1-yl)pyrimidine-2,4(1*H*,3*H*)-dione (9)

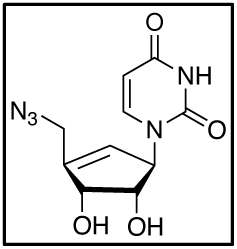

To intermediate **S2** (6 mg, 0.023 mmol, 1 equiv.) was added THF (250 μL) and concentrated HCl (25 μL). The reaction mixture was stirred at room temperature for 5 h. An excess amount of NaHCO_3_ was added to neutralize the reaction, followed by the addition of 1 mL MeOH. The resulting suspension was then filtered through a short Celite pad, rinsed with 2 × 2 MeOH, and concentrated. The resulting crude was resuspended with 1 mL H_2_O and subjected to HPLC purification with a linear gradient of 5-20% MeCN (0.1% TFA) in H_2_O (0.01% TFA) on a prep C18 column (Agilent 10 prep-C18 250 21.1 mm). Fractions containing desired products were then combined and lyophilized to afford the final product **9** (3.6 mg, 0.014 mmol, 59%) as white powder. 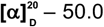 (*c* 0.36, MeOH); **IR** (film, cm^−1^): 3358 (br), 2921, 2104, 1683, 1467, 1393, 1261, 1205, 1116.^**1**^**H NMR** (400 MHz, MeOD) δ 7.39 (d, *J* = 8.0, 0.6 Hz, 1H), 5.81 (dd, *J* = 3.2, 1.9 Hz, 1H), 5.70 (d, *J* = 8.0 Hz, 1H), 5.52 – 5.40 (m, 1H), 4.52 (d, *J* = 5.8 Hz, 1H), 4.14 – 3.97 (m, 3H).^**13**^**C NMR** (100 MHz, MeOD) δ 166.4, 153.0, 146.7, 143.7, 128.5, 102.9, 77.9, 74.5, 67.9, 50.4. **HRMS** (ESI) *m/z* 288.0697 [(M+Na)^+^; calcd for C_10_H_11_N_5_O_4_Na^+^: 288.0703].

#### ((3a*S*,4*R*,6a*R*)-4-(3-benzoyl-2,4-dioxo-3,4-dihydropyrimidin-1(2*H*)-yl)-2,2-dimethyl-3a,6a-dihydro-4*H*-cyclopenta[*d*][1,3]dioxol-6-yl)methyl acetate (10)

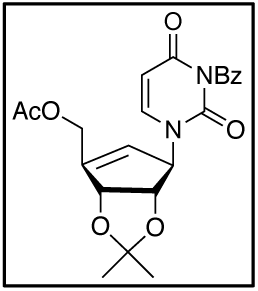

Triethylamine (66 uL, 0.48mmol, 3 equiv.) and acetyl chloride (AcCl) (17 μL, 0.24 mmol, 1.5 equiv.) were sequentially added to a solution of **5** (60 mg, 0.16 mmol, 1 equiv.) in CH_2_Cl_2_ (2 mL) at 0 °C. The reaction mixture was then warmed up to room temperature and stirred for another hour. A saturated solution of NH_4_Cl (2 mL) was added. The organic layer was extracted with 3 × 3 CH_2_Cl_2_, washed with H_2_O, and concentrated under reduced pressure. The crude material was purified through a 4g SiO_2_ flash cartridge with a linear gradient 65-95% EtOAc in hexanes to afford the intermediate **10** (56 mg, 0.13 mmol, 82%) as colorless oil. 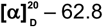 (*c* 1.3, CHCl_3_); **IR** (film, cm^−1^): 3080, 2988, 2935, 1742, 1703, 1663, 1599, 1440, 1372, 1236, 1179, 1236, 1058, 904, 731.^**1**^**H NMR** (400 MHz, CDCl_3_) δ 7.93 (d, *J* = 7.6 Hz, 2H), 7.78 – 7.61 (m, 1H), 7.50 (dd, *J* = 8.1, 7.7 Hz, 2H), 7.10 (d, *J* = 8.1 Hz, 1H), 5.81 (d, *J* = 8.1 Hz, 1H), 5.61 (brs, 1H), 5.34 (brs, 1H), 5.23 (d, *J* = 5.7 Hz, 1H), 4.79 (qt, *J* = 15.0, 1.8 Hz, 2H), 4.67 (d, *J* = 5.3 Hz, 1H), 2.12 (s, 3H), 1.42 (s, 3H), 1.34 (s, 3H). ^**13**^**C NMR** (100 MHz, CDCl_3_) δ 170.7, 168.7, 162.2, 149.7, 148.3, 141.2, 135.3, 131.5, 130.7, 129.3, 123.2, 113.1, 102.6, 84.1, 83.7, 68.6, 60.9, 27.3, 25.9, 20.9. **HRMS** (ESI) *m/z* 427.1508 [(M+H)^+^; calcd for C_22_H_23_N_2_O_7_+: 427.1500].

#### 1-((1*R*,4*R*,5*S*)-4,5-dihydroxy-3-methylcyclopent-2-en-1-yl)pyrimidine-2,4(1*H*,3*H*)-dione (11)

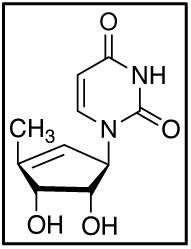

To a solution of the allylic acetate **10** (56 mg, 0.13 mmol, 1 equiv.) in 95% ethanol (4 ml) and cyclohexene (2 ml) was added 20 w% Pd(OH)_2_ on carbon (20mg, 1:3 catalyst substrate by weight). The resulting suspension was stirred under reflux overnight. The reaction mixture was then filtered, concentrated, and dried under high vacuum to afford 20 mg crude material as pale-yellow oil. In a similar manner as the preparation of **9**, 6 mg of the crude product was treated with 25 μL HCl in 250 μL THF to afford the final product **11** (8 mg, 0.037 mmol, 28% for two steps) as white solid. 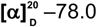 (*c* 0.24, MeOH); **IR** (film, cm^−1^): 3364 (br), 2921, 2851, 1680, 1468, 1393, 1251, 1203, 1116.^**1**^**H NMR** (400 MHz, MeOD) δ 7.39 (d, *J* = 8.0 Hz, 1H), 5.67 (d, *J* = 8.0 Hz, 1H), 5.52 – 5.35 (m, 2H), 4.37 (d, *J* = 5.8 Hz, 1H), 3.99 (dd, *J* = 5.9, 5.4 Hz, 1H), 1.88 (s, 3H).^**13**^**C NMR** (100 MHz, MeOH) δ 166.4, 153.1, 148.9, 143.5, 125.6, 102.7, 78.2, 77.2, 68.1, 15.1. **HRMS** (ESI) *m/z* 247.0685 [(M+Na)^+^; calcd for C_10_H_12_N_2_O_4_Na: 247.0689].

### Cloning, expression and purification of recombinant human UCK2 and CMPK1

Uridine cytidine kinase 2 (UCK2) was cloned into the pET-21a(+) vector using NdeI and NotI as restriction sites. This resulted in C-terminally 6×His-tagged UCK2. The resulting plasmid was transformed into *E. coli* BL21(DE3) cells by electroporation, recovered in SOC at 37 °C for 1 h and plated onto LB agar plates with carbenicillin overnight at 37° C. A single, isolated colony was inoculated into 30 mL LB supplemented with 50 µg/mL carbenicillin and grown at 37°C with shaking for 15 h. The next day, these cells were used to inoculate 1 L autoclaved LB with 50 µg/mL carbenicillin. The flask was shaken at 37 °C. When an optical density (OD_600_) of 0.7 was reached, protein expression was induced with 0.5 mM isopropyl β-D-1-thiogalactopyranoside (IPTG). The temperature was changed to 18 °C and flasks were shaken for an additional 15 h. The cell pellets were collected by spinning the media at 5000 rpm for 10 min. The pellets obtained were frozen by liquid nitrogen and stored in –80 °C for protein purification.

Cell pellets were thawed and resuspended in lysis buffer containing Tris (40 mM, pH 7.5), NaCl (10 mM), imidazole (10 mM), DTT (1 mM). Cells were lysed by sonication and centrifuged at 25,000g for 1 h. The supernatant was incubated with a slurry of Ni-NTA resin for 1h at 4 °C and loaded onto a column. The nickel column wash buffer was Tris (40 mM, pH 7.5), NaCl (10 mM), DTT (1 mM) and each wash step contained increasing concentrations of imidazole (10, 50, or 250 mM). After SDS-PAGE confirmation of the protein fractions, protein was further purified using fast protein liquid chromatography (FPLC) using buffer A (50 mM Tris-HCl pH 8, 1mM dithiothreitol (DTT), 10% glycerol) and elution buffer B (50 mM Tris-HCl pH 8, 1mM dithiothreitol (DTT), 10% glycerol with 500 mM NaCl) with changing gradient over a 20 minute from 2% to 100% Buffer B. FPLC eluents containing the protein was concentrated using Amicon centrifugal filter with 3 K cut-off and the enzyme was stored in storage buffer (50 mM Tris-HCl pH 7.5, 10% glycerol).

Human cytidine monophosphate kinase 1 (CMPK1) was purified as previously described (Deans et al., 2016).

### *In vitro* enzyme activity assays with recombinant human UCK2

To continuously monitor reaction progress spectrophotometrically, ATP hydrolysis was coupled to NADH oxidation via pyruvate kinase (PK) and lactate dehydrogenase (LDH) (Tomoike et al., 2017). Reactions were conducted at room temperature in 100 µL in 96-well plates (Greiner Bio-One, UV-Star, Half Area). Mixtures contained 20 mM HEPES (pH 7.2), 100 mM KCl, 2 mM MgCl_2_, 300 µM ATP, 0-500 µM uridine, 0-500 µM CPU analogs, 10 nM UCK2 (unless otherwise noted), 1 mM phosphoenolpyruvate, 500 µM NADH and 20 units/mL of PK and LDH. Progress was monitored in the linear region using a Biotek Synergy HT and kinetic and inhibition constants were determined using GraphPad Prism 7 (GraphPad Software).

### Enzymatic synthesis of CMP, CPU-MP, and 5-F-CPU-MP

Reactions were conducted at room temperature in 100 µL in Eppendorf tubes. Mixtures contained 20 mM HEPES (pH 7.2), 100 mM KCl, 2 mM MgCl_2_, 2.5 mM ATP, 5 mM substrates (cytidine, CPU and 5-F-CPU), and 2 µM UCK2. After gently mixing for 24 h on a rocking shaker, the reaction mixtures were heated for 3 minutes at 95 °C to denature UCK2 enzyme. Formation of CPUMP and 5-F-CPUMP were confirmed by LC-MS/MS (Figure S1).

### *In vitro* enzyme activity assays with recombinant human CMPK1

In a similar manner as UCK2 assay, ATP hydrolysis was coupled to NADH oxidation via pyruvate kinase (PK) and lactate dehydrogenase (LDH) to continuously monitor reaction progress spectrophotometrically. Reactions were conducted at room temperature in 50 µL in 96-well plates (Greiner Bio-One, UV-Star, Half Area). Mixtures contained 50 mM TrisHCl (pH 7.5), 50 mM KCl, 5 mM MgCl_2_, 2 mM DTT, 500 µM NADH, 1 mM PEP, 20 units/mL of PK and LDH. To 39.5 µL above reaction mixtures were add 5 µL 0-1 mM stock solutions of substrates (CMP from the commercial vendor and denatured UCK2 reaction mixtures containing newly synthesized CMP, CPU-MP and 5-F-CPUMP) in UCK2 assay buffer [20 mM HEPES (pH 7.2), 100 mM KCl, 2 mM MgCl_2_]. After the UV-readout at 340 nm reached stable in ca. 5 minutes, a mixture of ATP and CMPK1 in assay buffer [50 mM TrisHCl (pH 7.5), 50 mM KCl, 5 mM MgCl_2_, 2 mM DTT] was added to give a final reaction volume of 50 µL and a final ATP concentration of 500 µM and CMPK1 of 20-100 nM. Progress was monitored in the linear region using a Biotek Synergy HT, and kinetic constants were determined using GraphPad Prism 7 (GraphPad Software). Generation of CPU-DP and 5-F-CPU-DP was confirmed by LC-MS/MS (Figure S2).

### Sample preparation for LC-MS/MS analysis (Figure S3)

A549 cells were plated overnight at 80,000 cells/well in 24-well plates in complete DMEM. The next day, cells were treated with dipyridamole or 250 µM or 500 µM CPU, 5-F-CPU or 5’-F-CPU in DMEM additionally supplemented with 1 µM GSK983 and 5 µM uridine. Six or twelve hours after drug addition, the cell culture medium was removed and the whole plate was rapidly rinsed twice by dipping vertically into a beaker containing 37 °C Milli-Q water. The plate was then placed on dry ice, followed by the addition of 0.5 mL –20 °C lysis buffer (MeCN:MeOH:H_2_O = 2:2:1) containing 0.5 M formic acid and 450 nM uridine ^15^N_2_ monophosphate sodium salt (Sigma Aldrich). The solution was then sonicated for 3 × 5 minutes on ice for the metabolite extraction. Subsequently, the sample was frozen with liquid nitrogen, freeze-dried, re-suspended with 150 µL Milli-Q water, and finally filtered through a MultiScreen 96-well filter plate prior to LC-MS analysis.

### LC-MS/MS analysis of mono-, di- and triphosphates of cytidine, uridine, CPU and 5-F-CPU

LC-MS/MS was performed on an Agilent 1290 infinity II LC system tandem with 6470 triple quad mass spectrometer using ion pairing chromatography. LC separation was performed at 40 °C with a solvent flow rate of 0.25 mL/min on a Zorbax RRHD Extended-C18 column (2.1 × 150 mm, 1.8 µM). Buffer A contained 97% H_2_O, 3% MeOH, 5 mM tributylamine, 5.5 mM acetic acid and 1 µM medronic acid at pH=5.0;Buffer B contained ca. 100% MeOH, 5 mM tributylamine, 5.5 mM acetic acid and 10 µM medronic acid at pH=7.0 The initial mobile phase composition was 100% solvent A, and the gradient (%B) after sample injection was as follows: 0% at 2.5 min, 20% at 7.5 min, 45% at 14 min, 99% at 20 to 23 min, and 0% from 23.1 to 27.1 min. The autosampler temperature was set to 4 °C and 5 µL of sample was injected. Samples were measured in the negative ESI mode with capillary voltage at −3.5kV. Further source settings were as follows: gas temperature, 250 °C; gas flow, 13 L/min; nebulizer, 35 psi; sheath gas temperature, 325 °C; sheath gas flow, 12 L/min; nozzle voltage, 500 V; and delta EMV, – 200. Acquisition was performed in dynamic multiple reaction monitoring (dMRM) mode with fragmentation pattern and retention time setting as Table SI.

### DENV-2 luciferase reporter assays

A549 cells were plated overnight at 5,000 cells/well in 96-well plates in complete DMEM and incubated for 24h at 37°C. Cells were then treated with pyrimidine *de novo* synthesis inhibitors, and/or salvage inhibitors, and/or R1479 in DMEM supplemented with 20 µM uridine. Four hours after drug addition, cells were infected with DENV-2 virus at a MOI = 0.1. After 48 h or 72 h, DENV-Luc replication was monitored by the production of Renilla luciferase, which was measured using the Renilla-Glo Luciferase Assay System (Promega) according to the specifications of the manufacturer.

For the accompanying cell viability assay, A549 cells were seeded at 5,000 cells/well in 96-well plates in complete DMEM and incubated for 24h at 37°C. Cell were then treated with pyrimidine *de novo* synthesis inhibitors, and/or salvage inhibitors, and/or R1479 in DMEM supplemented with 20 µM uridine. Following 48 h or 72 h treatment, cell viability was monitored via ATP levels, which were measured using the CellTiter-Glo Luciferase Assay System (Promega) according to the specifications of the manufacturer.

For tests of the CPU-GSK983 combination (Figure 5B), A549 cells were plated overnight at 20,000 cells/well in 24-well plates in complete DMEM and incubated for 24 h at 37 °C. Cell were then treated with pyrimidine *de nov*o synthesis and/or salvage inhibitors in DMEM supplemented with 20 µM uridine. Four hours after drug addition, cells were infected with DENV-2 virus at a MOI = 0.01. After 48 h, DENV-Luc replication was monitored by the production of Renilla luciferase, which was measured using the Renilla-Glo Luciferase Assay System (Promega) according to the specifications of the manufacturer.

For tests of the CPU-GSK983 combination (Figure 5B),For the accompanying cell viability assay, A549 cells were seeded into 24-well plates at a density of 20,000 cells/well incubated for 24 h at 37 °C. Cell were then treated with pyrimidine *de nov*o synthesis and/or salvage inhibitors in DMEM supplemented with 20 µM uridine. Following 48h treatment, cells were harvested, and the density of viable cells was determined by flow cytometry (FSC/SSC) using a BD Accuri C6 Flow Cytometer.

### Replicon assays

Transfected A549 cells were plated at 15,000 cells/well in 96-well plates, and immediately treated with R1479, and/or, pyrimidine *de novo* synthesis inhibitors, and/or salvage inhibitors in complete DMEM medium supplemented with 20 µM uridine without antibiotics (Marceau et al, 2016). Forty-eight hours after drug treatment, cells were lysed and subjected to Renilla luciferase detection, which was measured using the Renilla-Glo Luciferase Assay System (Promega) according to the specifications of the manufacturer.

## SUPPLEMENTAL FIGURES

**Figure S1.**
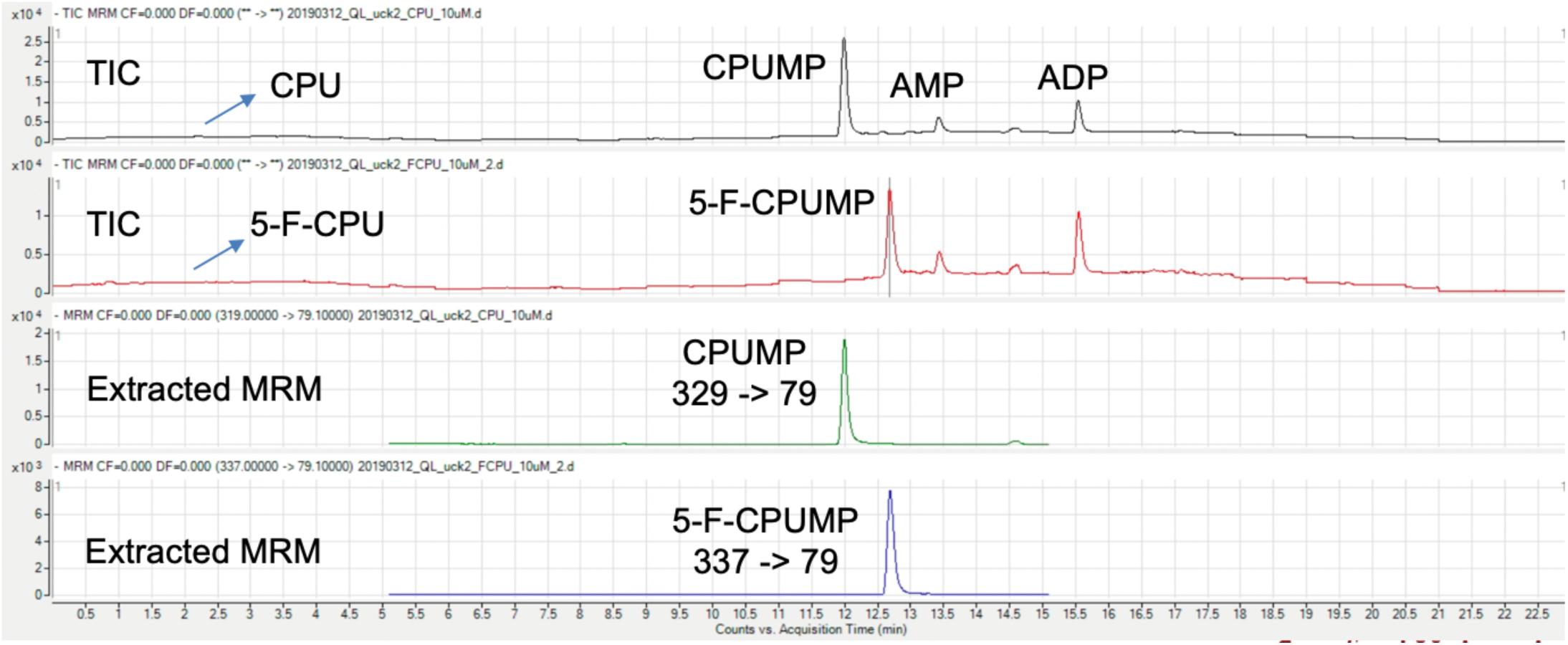
Identities of CPU-MP and 5-F-CPU-MP were confirmed by LC-MS/MS, related to Figure 3*G*. TIC: Total Ion Chromatogram; MRM: Multiple Reaction Monitoring.

**Figure S2.**
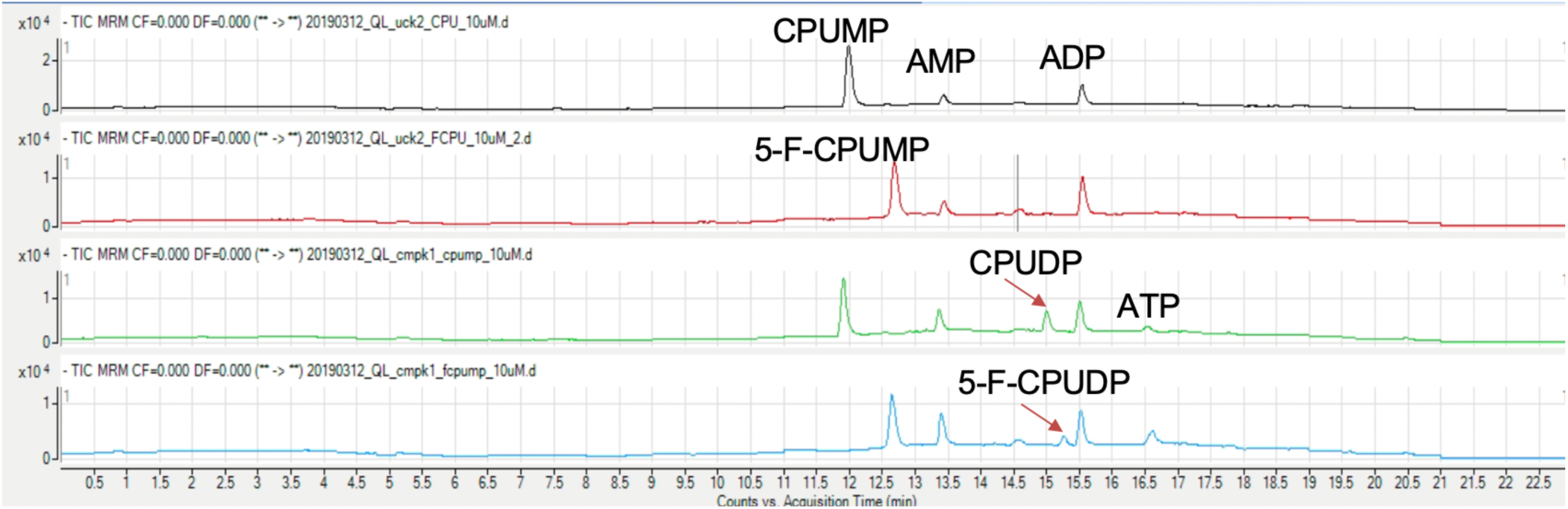
LC-MS/MS monitoring of CMPK1-catalyzed conversion of CPU-MP and 5-F-CPU-MP to the corresponding diphosphates, related to Figure 3*G*.

**Figure S3.**
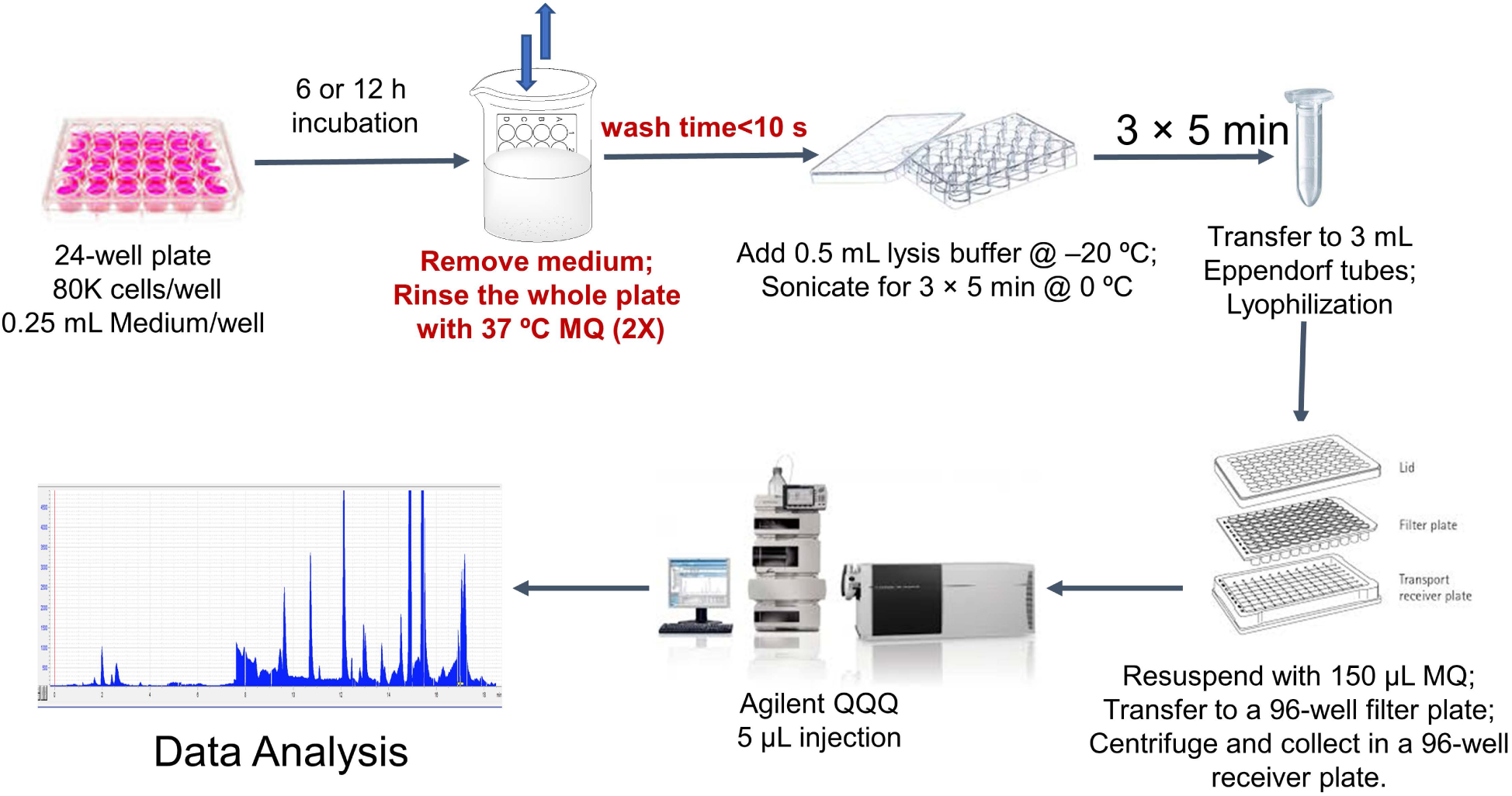
Flow chart of cell sample preparation for metabolite analysis by LC-MS/MS.

**Figure S4:**
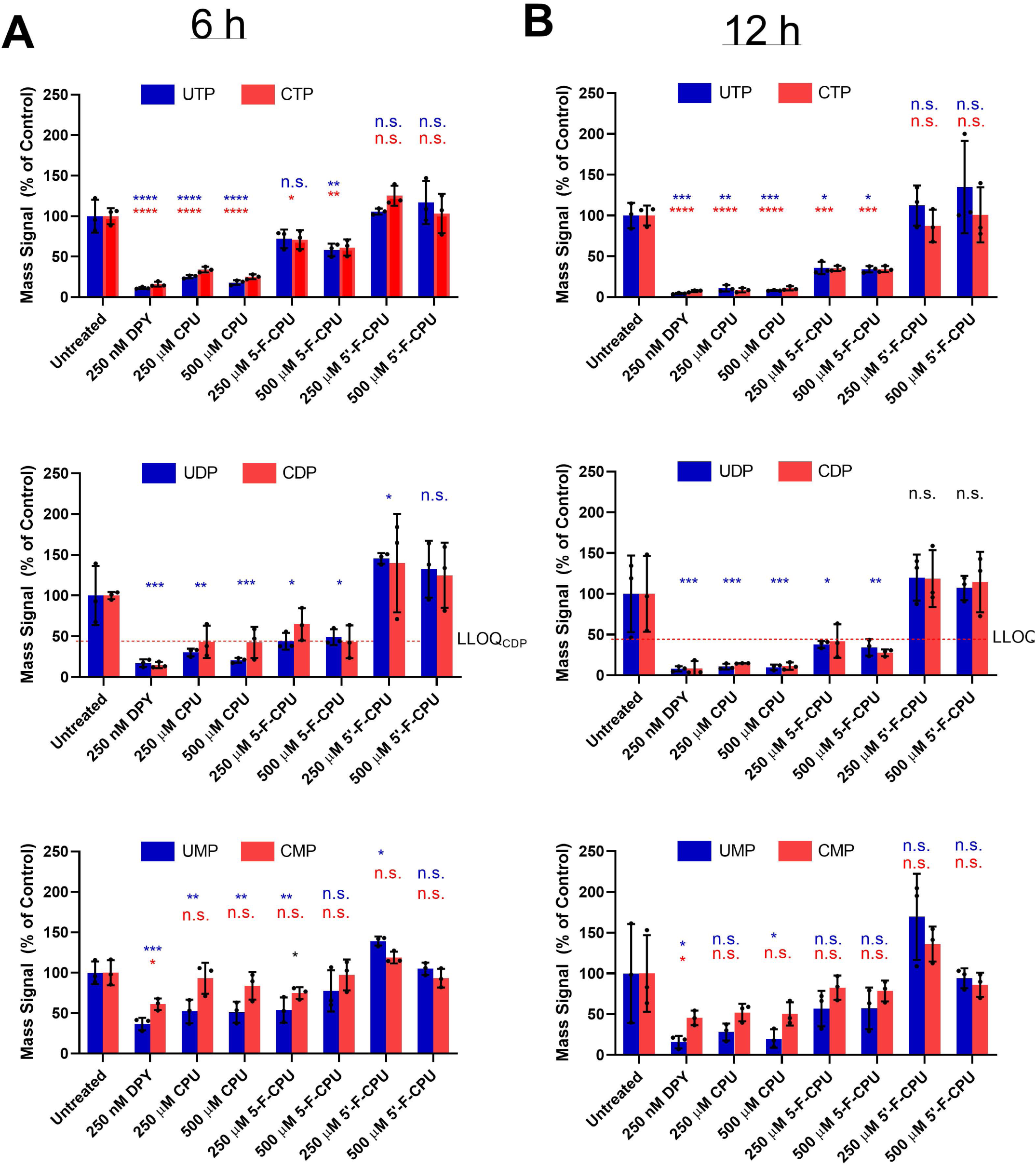
Analysis of intracellular pyrimidine pools in A549 cells upon drug addition. *(A)* LC-MS analysis of intracellular pyrimidine TP, DP and MP pools after 6 h indicated drug treatment. Row 1 data is repeated from Figure 4*A*. *(B)* LC-MS analysis of intracellular pyrimidine TP, DP and MP pools after 12 h indicated drug treatment. In all assays, cell culture medium was supplemented with 5 µM uridine and 1 µM GSK983. Dipyridamole (DPY)—a nucleoside transport inhibitor—was used as a positive control. Bars indicate means ± S.D, n = 3 and dots represent individual values. The statistical significance of comparisons to the untreated controls are indicated for uridine compounds (blue) and cytidine compounds (red) with * (p<0.05), ** (p<0.01), *** (p<0.001) or n.s. (p>0.05) using one-way ANOVA, Dunnett’s test. In the DP graphs, the red line indicates the lower limit of reliable linear quantitation for CDP. Statistical tests were not carried out for CDP as many samples fell below the LLOQ.

**Figure S5:**
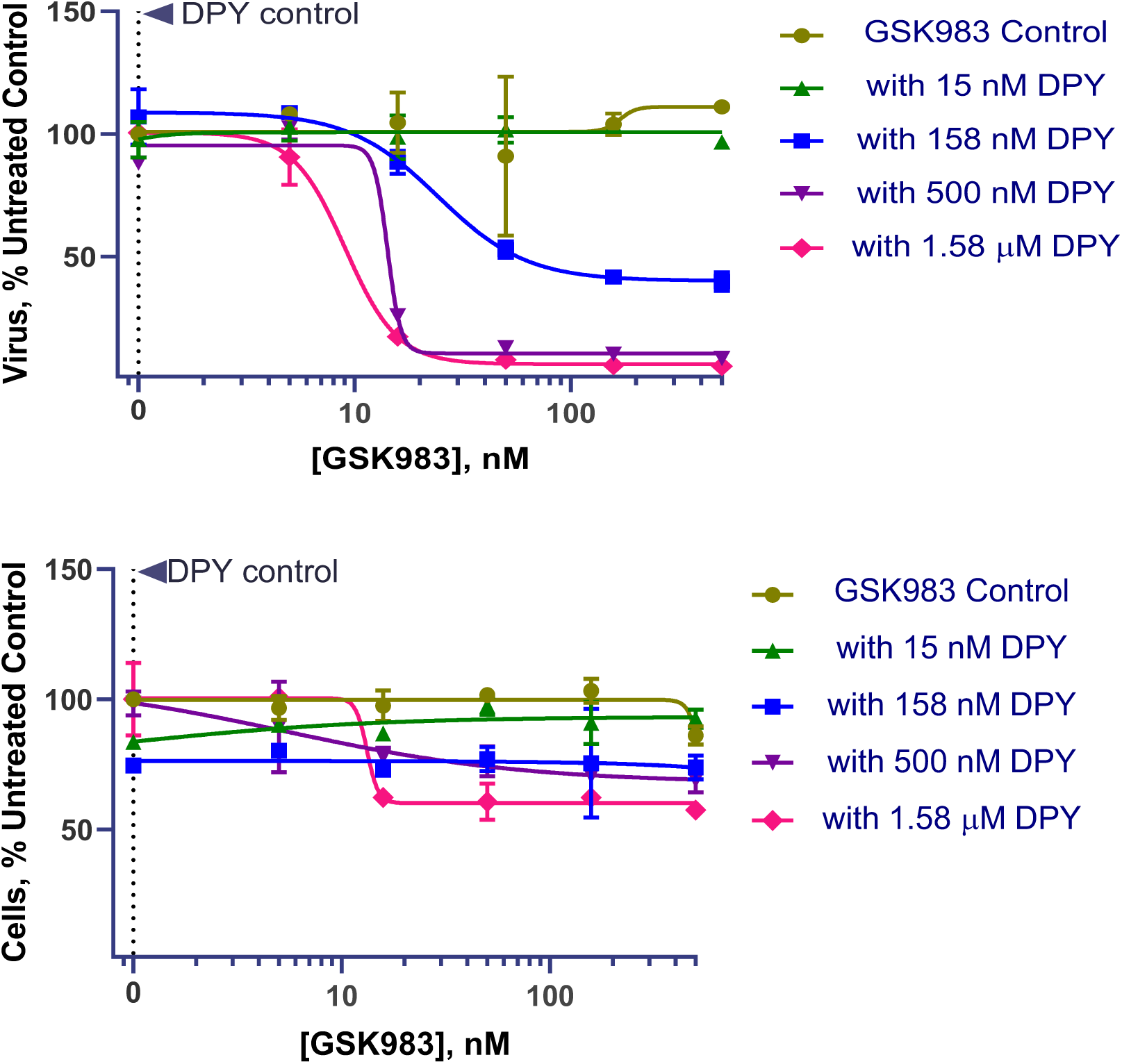
Effects of DPY-GSK983 combination therapy on dengue virus replication and cell proliferation in the presence of exogenous uridine at 48 h. Error bars represent ± S.D. of two replicates.

**Figure S6.**
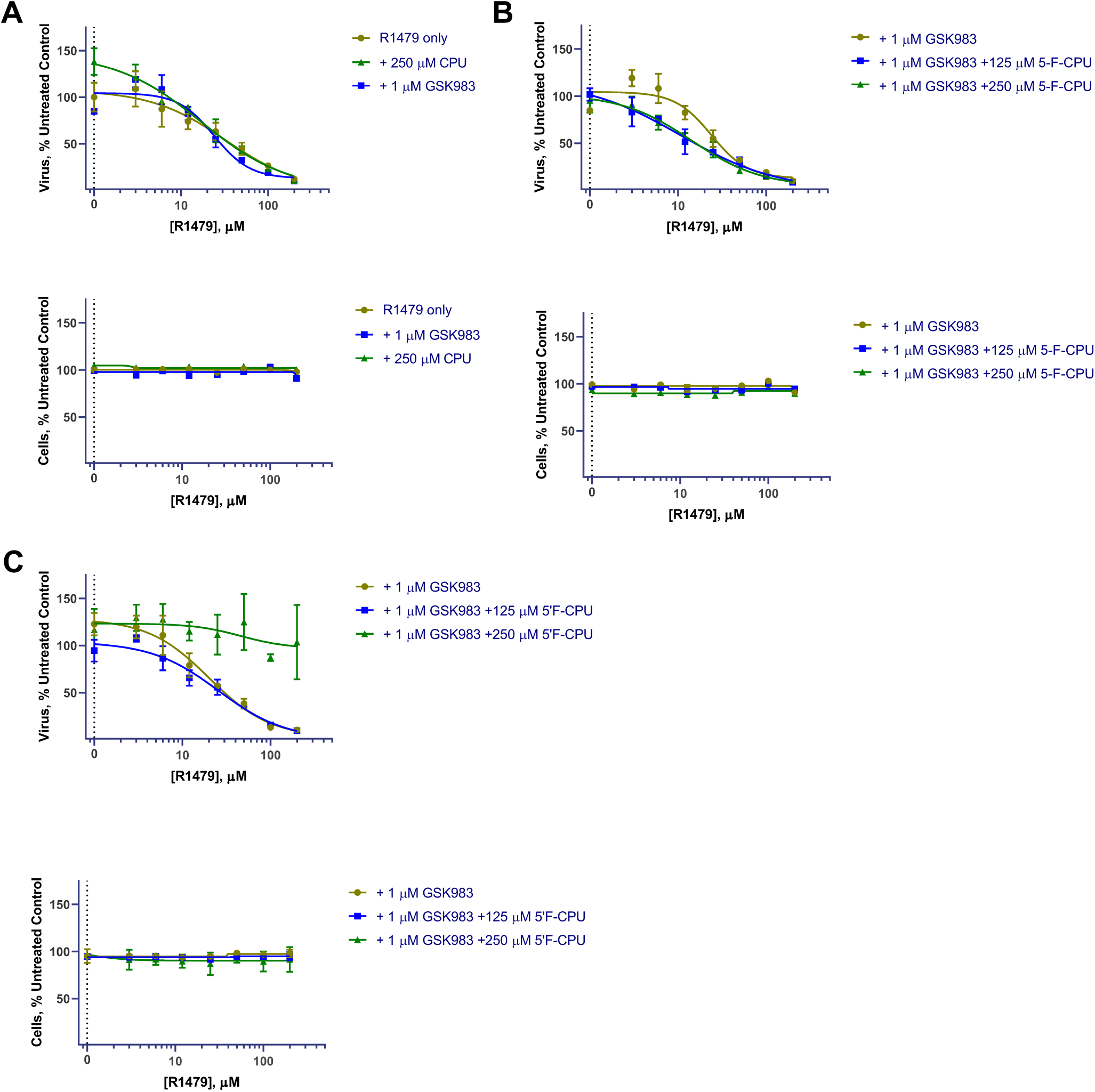
Potentiating the effect of R1479 on dengue virus replication in the presence of exogenous uridine at 48 h. *(A).* Luminescence of DENV-2 infectious clone and CellTiterGlo viability reagent in A549 cells upon treatment with R1479, R1479 + 250 µM CPU, or R1479 + 1 µM GSK983. Data is expressed as % of untreated no drug control. *(B).* Effect of 5-F-CPU on dengue virus infection and cell viability. *(C)* Effect of 5’-F-CPU on dengue virus infection and cell viability. In all assays, the culture medium was supplemented with 20 µM uridine and as noted, with 1 µM GSK983 to block *de novo* pyrimidine biosynthesis.

**Figure S7:**
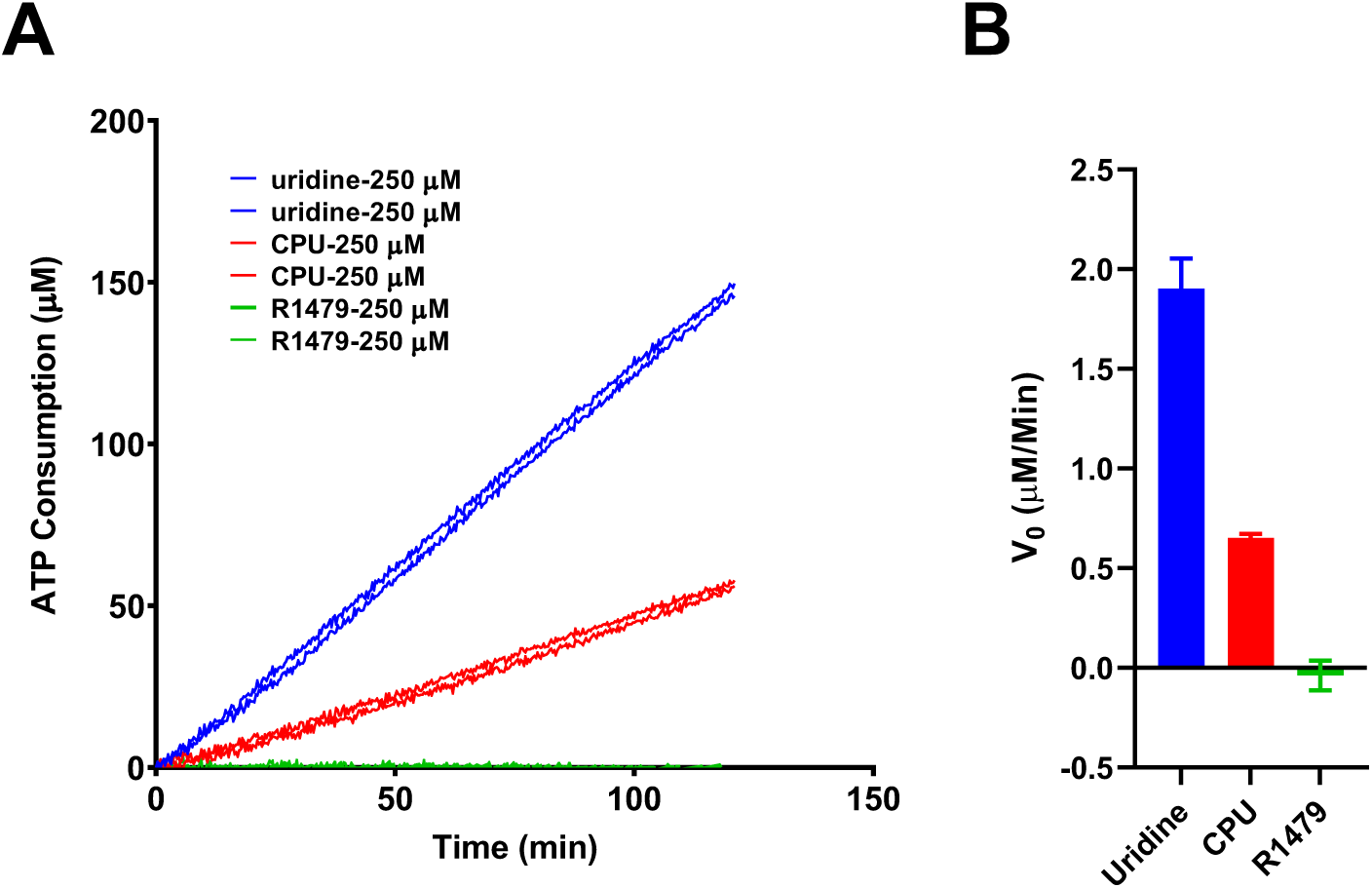
R1479 (4’ azido-cytidine) is not phosphorylated by UCK2. *(A)* ATP consumption in duplicate reactions containing 3 nM UCK2 and 500 uM ATP with 250 µM uridine, CPU or R1479 respectively. *(B)* Initial rates for three substrates. Error bars represent ± S.D. of technical duplicates.

**Table S1.**
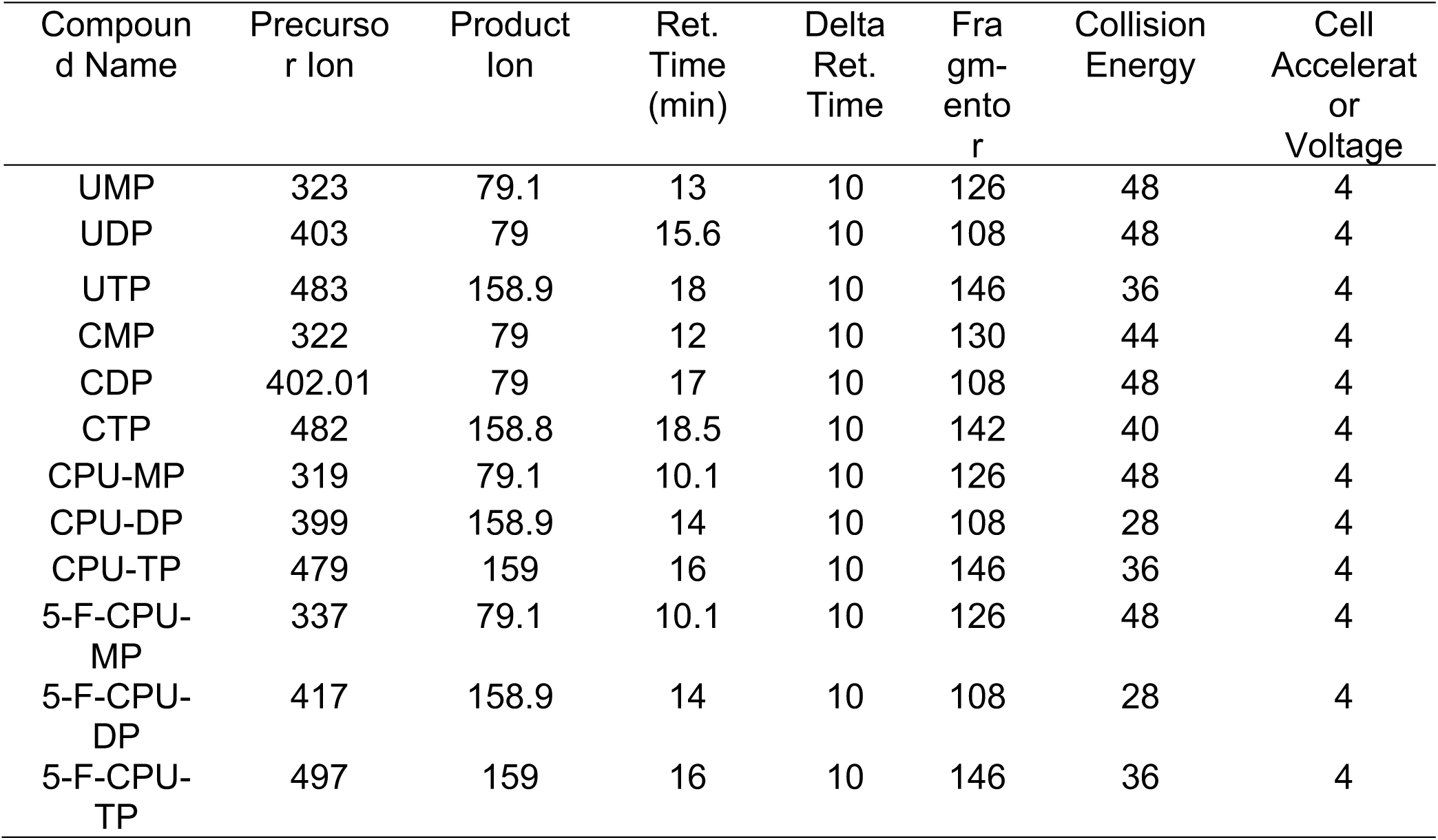
Retention time and products monitored by LC-MS.

| Compound Name | Precursor Ion | Product Ion | Ret. Time (min) | Delta Ret. Time | Fragmentor | Collision Energy | Cell Accelerator Voltage |
| --- | --- | --- | --- | --- | --- | --- | --- |
| UMP | 323 | 79.1 | 13 | 10 | 126 | 48 | 4 |
| UDP | 403 | 79 | 15.6 | 10 | 108 | 48 | 4 |
| UTP | 483 | 158.9 | 18 | 10 | 146 | 36 | 4 |
| CMP | 322 | 79 | 12 | 10 | 130 | 44 | 4 |
| CDP | 402.01 | 79 | 17 | 10 | 108 | 48 | 4 |
| CTP | 482 | 158.8 | 18.5 | 10 | 142 | 40 | 4 |
| CPU-MP | 319 | 79.1 | 10.1 | 10 | 126 | 48 | 4 |
| CPU-DP | 399 | 158.9 | 14 | 10 | 108 | 28 | 4 |
| CPU-TP | 479 | 159 | 16 | 10 | 146 | 36 | 4 |
| 5-F-CPU-MP | 337 | 79.1 | 10.1 | 10 | 126 | 48 | 4 |
| 5-F-CPU-DP | 417 | 158.9 | 14 | 10 | 108 | 28 | 4 |
| 5-F-CPU-TP | 497 | 159 | 16 | 10 | 146 | 36 | 4 |

